# Liver regeneration by a population of midzone-located mesenchymal-hepatocyte hybrid cells

**DOI:** 10.1101/2023.10.16.562602

**Authors:** Guo Yu, Shaoyang Zhang, Ana Romo, Soma Biswas, Baojie Li, Jing Li

## Abstract

Regenerative medicine relies on deep understanding of the mechanisms of organ repair and regeneration. The liver, an organ with critical metabolic functions carried out by hepatocytes located in zones 1-3 of liver lobules, has the capacity to fully regenerate itself, which is mainly attributable to midzone hepatocytes. Yet, how differentiated midzone hepatocytes execute transzone regeneration and quickly recover most of the liver mass remains a myth. Here, we uncover a mesenchymal-hepatocyte population (13.7% of total hepatocytes) that are derived from Twist2-lineage EpCAM^+^ progenitors, midzone-located, highly polyploidy, and equipped with great mitogenic and migratory capabilities to the detriment of metabolism. They regenerate about 50% of new hepatocytes and repopulate zones 1 and 3 in liver regeneration. Mechanistically, expansion of these cells is negatively controlled by Notch1 signaling. This study has thus uncovered a hepatocyte subpopulation with great proliferation potential and important mechanisms of liver regeneration.

## Introduction

Hepatocytes originate from bipotential hepatoblast progenitor cells during embryonic development under the control of signaling molecules including Wnt, Notch, HGF, and BMP(Pepe-Mooney et al., 2019; Russell & Monga, 2018; Segal et al., 2019; Siebel & Lendahl, 2017). They undergo zonation along the porto-central axis of the liver lobules, with different zones executing distinct metabolic functions(Aizarani et al., 2019; Halpern et al., 2017). For example, zone 1 (periportal) hepatocytes carry out mainly β-oxidation and gluconeogenesis; zone 3 (pericentral) hepatocytes carry out mainly glycolysis, lipogenesis, and drug detoxification; and zone 2 (midzone) hepatocytes show mixed activities(Ben-Moshe et al., 2019).

The liver is constantly exposed to viruses, bacteria, alcohol, and toxic agents brought in by nutrient-carrying blood from the gastrointestinal tracts, which damage the liver and increase the risk of hepatitis, cirrhosis, or liver cancer(Banales et al., 2019; Ramachandran et al., 2019; Romeo, 2019). In spite of slow hepatocyte turnover, mammalian livers have enormous regeneration capacities, for example, human liver can be fully regenerated within 4 months after 3/4 of the liver is removed(Bangru & Kalsotra, 2020). While stem/progenitor cells marked by Axin, Sox9, or Tert or generated from dedifferentiation of hepatocytes or cholangiocytes were initially suggested to drive liver homeostasis and regeneration(Ang et al., 2019; Font-Burgada et al., 2015; Lin et al., 2018; Miyajima et al., 2014; Pu et al., 2016; Raven et al., 2017; Sato et al., 2019; Wang et al., 2015), later studies showed that all hepatocytes undergo hypertrophic growth and hyperplasia, generating regenerative patches that help repair the liver stochastically and locally(Chen et al., 2020; Matsumoto et al., 2020; Sun et al., 2020). If all the hepatocytes are participating, they would need to divide only 2 times to recover the full liver mass in partial (2/3) hepatectomy (PHx) mouse models(Monga, 2020).

However, recent genetic studies show that liver homeostasis and regeneration are mainly executed by midzone hepatocytes(He et al., 2021; Wei et al., 2021). Most of the liver injuries occur to zone 1 or 3 and it is likely that zone 2 hepatocytes are protected from the damage (Andersson, 2021; Forbes & Newsome, 2016). However, to accomplish the regeneration task, midzone hepatocytes need not only to undergo massive expansion but also to repopulate zones 1 and 3. This presents a challenge as hepatocytes, the differentiated epithelial cells, in general have limited proliferation and migration capacities.

The liver also contains hepatic stellate cells (HSCs) and fibroblasts (designated mesenchymal cells (MCs) here)(Lua et al., 2014; Ramachandran et al., 2019), whose main function is secretion of extracellular matrix (ECM). Here, our lineage tracing experiments using 5 Cre lines driven by mesenchymal stromal cell (MSC) markers identified a mesenchymal-hepatocyte hybrid population (13.7% of total hepatocytes) that express signature genes of both lineages. They appear to be derived from Twist2-expressing hepatomesenchyme of early embryos and are mainly located in the midzone and highly polyploidy and display great proliferative and migratory abilities at the sacrifice of metabolism. During liver regeneration, these cells display proliferation rate 2-3 times greater than conventional hepatocytes and replenish zones 1 and 3, accompanied by conversion from polyploids to diploids. Overall, this small population of cells could regenerate about 50% of the new hepatocytes. Our gene ablation and pharmacological studies also elucidated an important mechanism by which expansion of these hybrid cells is controlled. Collectively, these findings identify a midzone-located mesenchymal-hepatocyte population as a major driving force of liver regeneration.

## Results

### Mesenchymal marker Twist2 labels a hepatocyte subpopulation

To understand the ontology of liver mesenchymal cells (MCs)(Lua et al., 2014), we crossed several Cre mouse lines, which are used to study bone marrow mesenchymal stromal cells (BM-MSCs), to *Rosa-tdTomato* reporter mice to fate map these lineages in the liver. We found that Col1α2, αSMA. Gli1, and Vimentin marked MCs, while surprisingly, Twist2 marked both MCs and a small portion of hepatocytes (Figure 1A). Twist2 is a helix-loop-helix transcription factor and *Twist2-Cre* mice are widely used to study skeletal development(Yu et al., 2003). Immunostaining showed that liver Tomato^+^ cells in *Tomato^Twist2^* mice were negative for CD31 (a blood vessel marker), CD45 (a Kupffer cell marker), and K19 (a bile duct cell marker) (Figure 1B). However, Tomato^+^ cells with hepatocyte morphology expressed hepatocyte marker Hnf4α; and 99% of Tomato^+^ non-hepatocyte cells expressed mesenchymal marker Vimentin and 26% expressed PDGFRα (Figure 1B and Figure S1A). On average, 36.4% of the liver MCs were Tomato^+^ in *Tomato^Twist2^* mice (Figure 1C). These results suggest that Twist2 genetically marks subpopulations of hepatocytes and MCs in mouse liver.

**Fig. 1.**
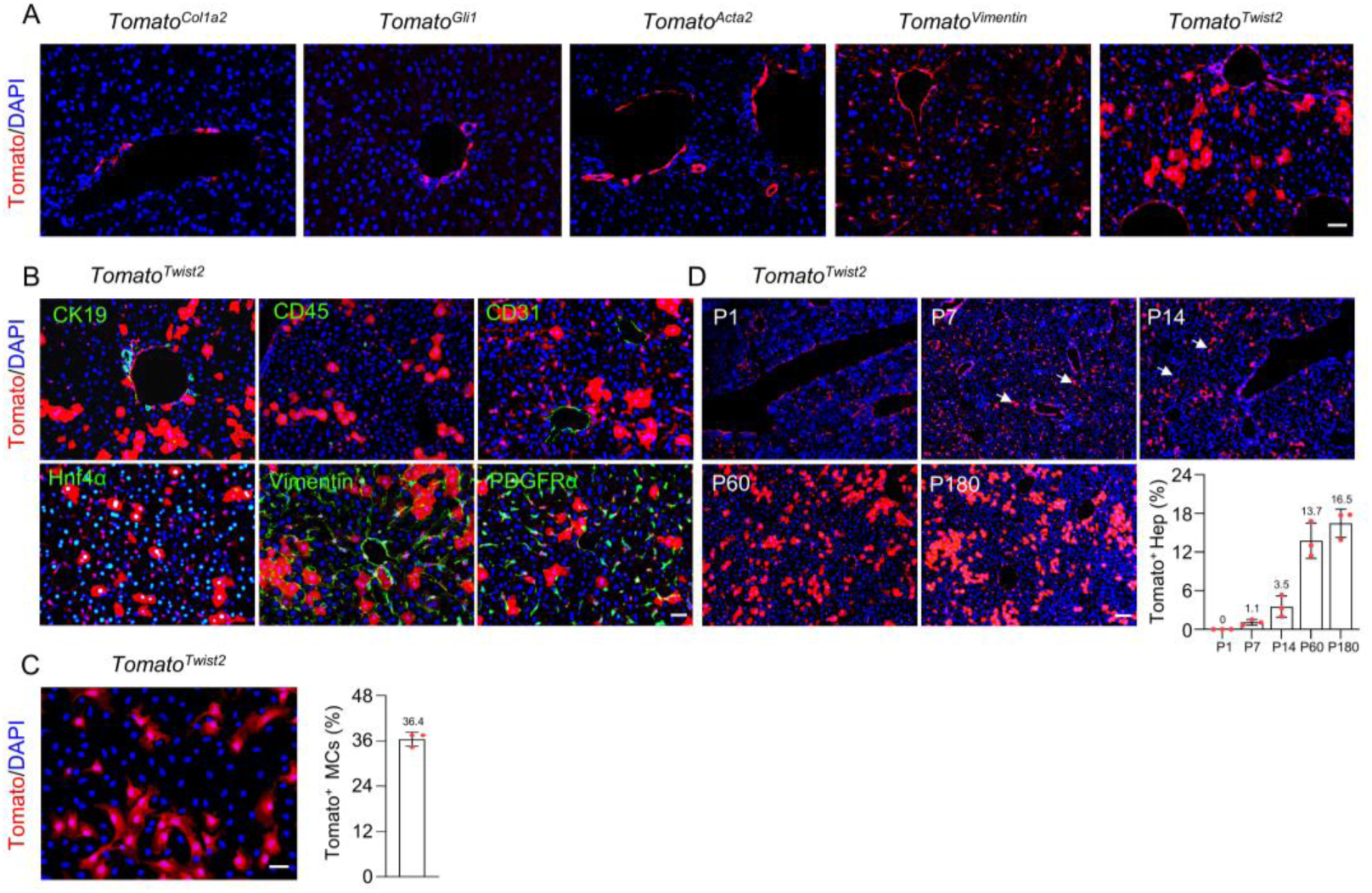
Twist2-lineage mesenchymal-hepatocytes are generated postnatally. **(A).** Results of lineage tracing using *Col1a2-CreERT*, *Gli1-CreERT*, *Acta2-Cre, Vim-CreERT*, and *Twist2-Cre* mouse lines crossed with the *Rosa-tdTomato* reporter line. Male mice over 2 months of age were used. For *CreERT* lines, TAM was injected at 1 month of age, and the mice were sacrificed 1.5 months later. Scale bar, 50 µm. **(B).** Illustrative immunostaining results of liver sections of adult *tdTomato^Twist2^* mice for CD31, CD45, K19, Vimentin, PDGFRα, or Hnf4α. Scale bar, 50 µm. **(C).** The percentages of Tomato^+^ cells in MCs cultures. Liver MCs were prepared from adult *Tomato^Twist2^*mice and plated and cultured overnight. Tomato^+^ cells were counted (6 views) under microscope and normalized to the number of DAPI^+^ cells. n=3. Scale bar, 50 µm. **(D).** Tracing of Twist2-lineage hepatocytes in *Tomato^Twist2^* mice of various ages. Right panel: quantitation data. n=3 for each age group. Scale bar, 100 µm.

### Ablation of one *Twist2* allele in *Twist2-Cre* mice does not affect liver development or regeneration

In *Twist2-Cre* mice, one allele of *Twist2* is disrupted due to insertion of the *Cre* cassette into the coding region(Yu et al., 2003). However, *Twist2-Cre* mice showed a normal liver weight-to-body weight (LW/BW) ratio and liver structure (Figure S1B,C). Immunostaining revealed no alteration in the numbers of Vimentin^+^ MCs or PCNA^+^ proliferating cells or activation of pro-proliferating pathways including ERK and mTOR in the liver of *Twist2-Cre* mice (Figure S1D). Moreover, the mice showed normal regeneration processes upon CCl_4_-induced liver injury or partial (2/3) hepatectomy (PHx) (Figure S1E). Although Twist2 has been shown to promote or inhibit cell proliferation in different contexts(Zhang et al., 2015), we show here that deletion of one allele of *Twist2* does not affect liver development, growth, or regeneration, suggesting that the *Twist2-Cre* mouse line can be safely used to study liver development or regeneration.

### Twist2-lineage hepatocytes are derived from EpCAM^+^ progenitors postnatally

Analysis of *Tomato^Twist2^* mice of various ages revealed that Twist2 mainly marked non-hepatocyte cells in the liver at postnatal day 1 (P1) (Figure 1D), while Tomato^+^ hepatocytes started to emerge at P7, peaked at P60, but did not significantly expand further in adult mice (Figure 1D and later results). The average of Tomato^+^ hepatocytes, based on 10 adult mice, was 13.7%.

To further understand the Twist2-lineage cells, we performed scRNA-seq analysis of Tomato^+^ cells isolated from the livers of P1 or P14 *Tomato^Twist2^* mice. We sequenced 10000 cells at a depth of 2500 genes per cell in each sample. t-SNE analyses revealed 3 major Tomato^+^ subgroups (Figure 2A-C): hepatocytes, MCs, and EpCAM^+^ cells. The EpCAM^+^ cells could be further classified into EpCAM^high^, EpCAM^med^, and EpCAM^low^ subpopulations, which showed similar gene expression profiles (Figure 2C). In addition, a population of Tomato^+^ cells expressing proerythroblast genes was detected in P1 but not P14 mice (Figure S2A-C), which might represent transit cells in fetal/neonatal livers and were not further analyzed. In addition, the EpCAM^med^ and EpCAM^low^ subgroups were not observed in P14 mice, which only contained the EpCAM^high^ subgroup (Figure 2D). Surprisingly, Twist2 is only expressed in some MCs but not in the EpCAM^+^ subgroups in P1 or P14 mice (Figure S2D), suggesting that EpCAM^+^ cells are derived from Twist2-expressing cells during early development, which have lost *Twist2* expression later.

**Fig. 2.**
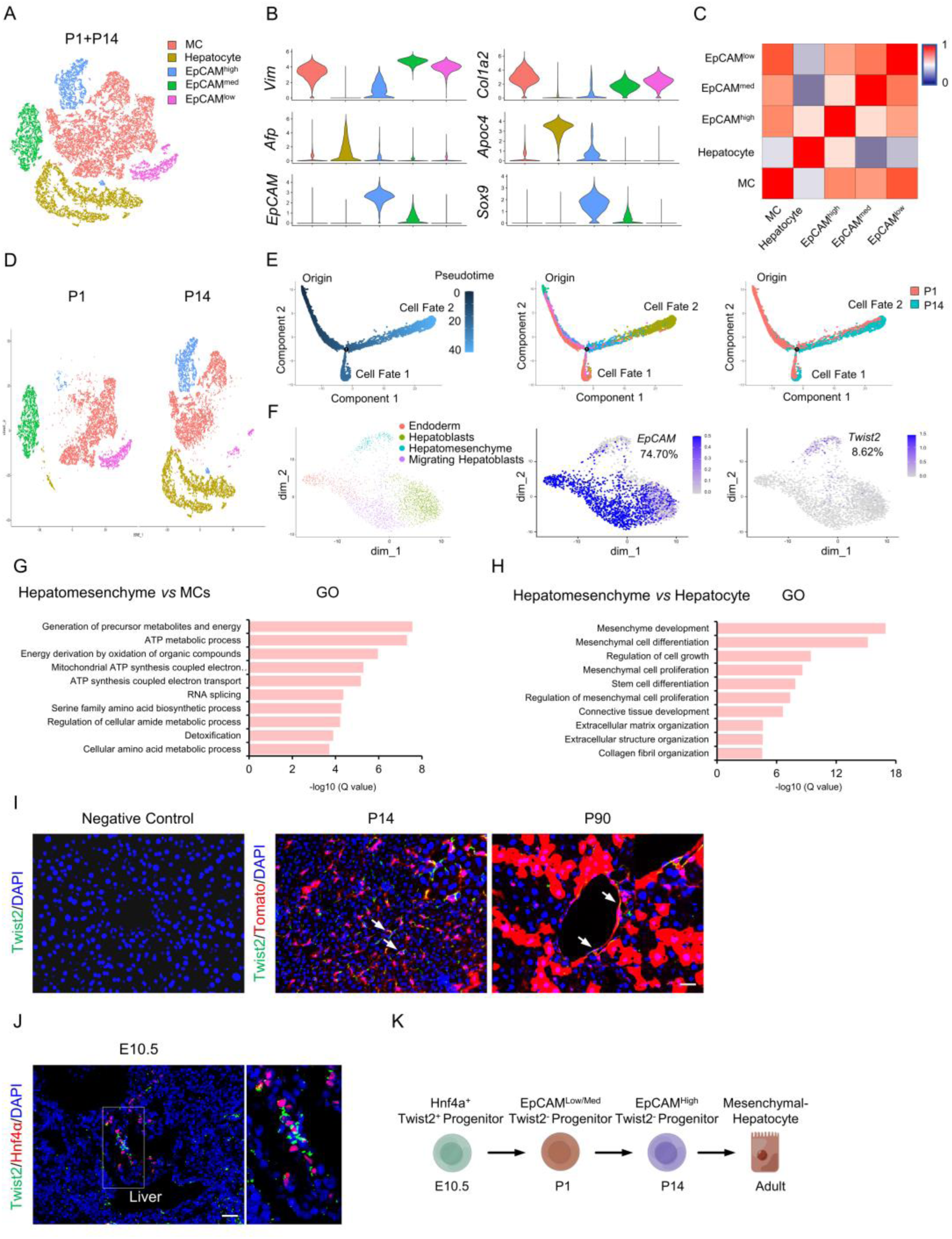
EpCAM^+^ progenitors generate Twist2-lineage hepatocytes and MCs. **(A).** t-SNE analysis of Tomato^+^ liver cells (combined cells from P1 and P14 *Tomato^Twist2^* mice). **(B).** Violin plots showing expression of marker genes for MCs, hepatocytes and EpCAM^+^ cells. **(C).** Heatmap analysis of the top 200 genes expressed in various Tomato^+^ subgroups. **(D).** Comparison of Tomato^+^ subgroups from the livers of P1 and P14 mice. **(E).** Trajectory analysis of Twist2-lineage subpopulations. **(F).** t-SNE analysis of EpCAM and Twist2 expression from public scRNA-seq datasets. **(G).** GO analysis of hepatomesenchyme compared to MCs. **(H).** GO analysis of hepatomesenchyme compared to hepatocytes. **(I).** Illustrative immunostaining results for Twist2 on liver sections of P14 and P90 *Tomato^Twist2^* mice. Scale bar, 50 µm. **(J).** Illustrative immunostaining results for Twist2 and Hnf4α on sections of E10.5 embryos. Scale bar, 50 µm. **(K).** A model explaining how early progenitors generate Twist2 lineage mesenchymal-hepatocytes in mice.

While in P1 pups, few of Tomato^+^ cells were hepatocytes, approximately 40% of the Tomato^+^ cells were hepatocytes in P14 mice (Figure S2B). Trajectory analysis revealed that cells in the EpCAM^med^ subgroup might first convert to EpCAM^high^ cells and then to hepatocytes (Figure 2E). Comparison of lineage signature gene expression revealed that EpCAM^med^ and EpCAM^low^ cells also expressed mesenchymal markers but not immune or endothelial cell markers (Figure S2E-G). Moreover, EpCAM^low^ cells showed a gene expression pattern close to MCs, compared to EpCAM^high^ or EpCAM^med^ cells (Figure 2C and Figure S2G). These findings suggest that EpCAM^low^ progenitor cells give rise to hepatocytes and MCs.

Gene ontology (GO) analysis confirmed that Twist2 lineage hepatocytes were enriched in metabolism genes whereas MCs were enriched in ECM genes (Figure S3A, B). Interestingly, EpCAM^med^ cells showed enriched expression of genes related to proliferation and macromolecule synthesis, EpCAM^high^ cells were enriched in genes related to proliferation and epithelial development, and EpCAM^low^ cells expressed some ECM genes (Figure S3C-E), consistent with the trajectory analysis results that the EpCAM^med^ subgroup might first convert to EpCAM^high^ cells and then to hepatocytes (Figure 2E).

### Detection of Twist2-expressing progenitors in E10.5 liver

Our scRNA-seq data suggest that *Twist2* is expressed in MCs but not EpCAM^+^ progenitors in P1 or P14 livers (Figure S2D). Analysis of scRNA-seq data from public datasets (GSE125688 and GSE137720)(Dobie et al., 2019; Pepe-Mooney et al., 2019) also revealed that Twist2 is expressed in a small population of MCs but not hepatocytes (clustered based on *Albumin* expression levels) in adult mice and that Twist2 expression is not altered during liver regeneration (Figure S4A,B). These results suggest there might exist a Twist2-expressing progenitors in early embryonic liver, which give rise to EpCAM^+^ cells in newborn mice and then Twist2 lineage hepatocytes in adult mice.

Indeed, a recent scRNA-seq study has uncovered a *Twist2-*expressing hybrid hepatomesenchymal cell type at E10.5(Lotto et al., 2020), whose daughter cells and function have not been investigated. We re-analyzed the scRNA-seq data and found that 8.62% of hepatocytes expressed Twist2, a portion of which also expressed EpCAM (Figure 2F). GO analyses revealed that the *Twist2-*expressing hepatomesenchymal cells expressed both stromal and hepatocyte signature genes (Figure 2G, H).

To confirm the above finding, we immunostained liver sections of *Tomato^Twist2^*mice of various ages for Twist2 expression. Although Twist2 is a transcription factor, it is also detected in the cytoplasm(Liu et al., 2017). We found that Twist2 was detectable in Tomato^+^ MCs but not Tomato^+^ hepatocytes in P14 or P90 mice (Figure 2I), consistent with scRNA-seq results. However, in E10.5 embryos, some of the Hnf4α^+^ cells were positive for Twist2 in the liver anlage region (Figure 2J). Both the scRNA-seq and immunostaining data suggest that Twist2 lineage hepatocytes are derived from Twist2^+^ progenitors during early embryonic development via Twist2^-^EpCAM^+^ intermediates (Figure 2K).

### Midzone location and polyploidy of Twist2-lineage hepatocytes

Most Tomato^+^ hepatocytes were located in the midzone, with some at the periportal region but few at the pericentral region (Figure 3A). Similar results were obtained from tracing experiments with *mTmG^Twist2^* mice, in which Twist2-expressing cells and their progenies were GFP^+^, while all other cells were Tomato^+^ (Figure 3B). This was also confirmed by immunostaining for pericentral zone hepatocyte marker CYP2e1 and periportal zone hepatocyte marker CDH1 (Figure 3C)(Ben-Moshe et al., 2019). Genetic studies have identified a few zone-specific markers(Wei et al., 2021), and analysis of our scRNA-seq data revealed that Twist2-lineage hepatocytes expressed *Cyp1a2*, *Gls2*, and *Ccnd1*, which mark portions of midzone hepatocytes, but not zone 3-specific marker G/S (Figure S5). Twist2-lineage hepatocytes did not express *Hamp2*, a marker for some midzone hepatocytes(Halpern et al., 2017), suggesting that Twist2 and Hamp2 mark different subpopulations of the midzone hepatocytes.

**Fig. 3.**
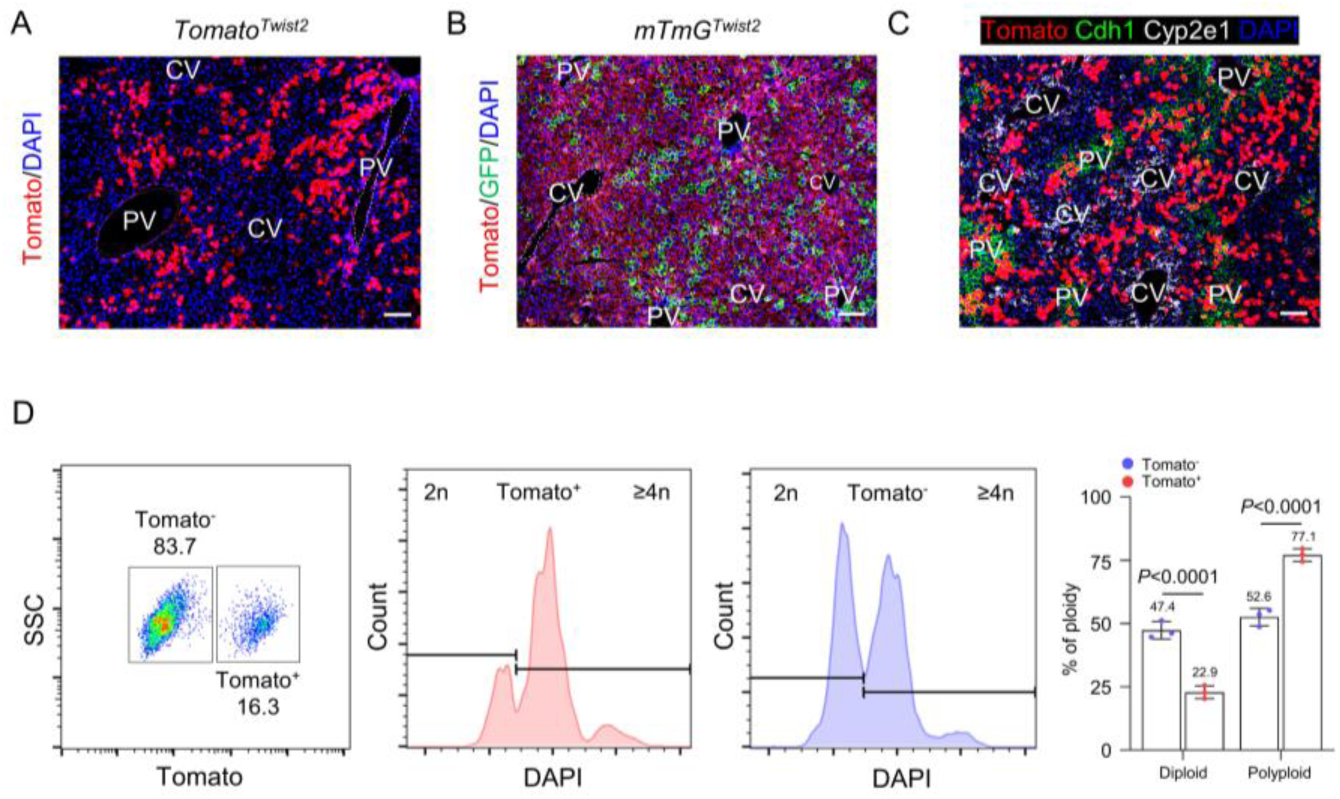
Midzone location and polyploidy features of mesenchymal-hepatocytes. **(A).** Location of Tomato^+^ hepatocytes in the liver of adult *Tomato^Twist2^* mice. Scale bar, 100 µm. **(B).** Illustrative image showing the tracing results of adult male *mTmG^Twist2^* mice. Scale bar, 100 µm. **(C).** Illustrative immunostaining results for CYP2e1 (zone 3) and CDH1 (zone 1) in liver sections from adult *Tomato^Twist2^* mice. Scale bar, 100 µm. **(D).** Flow cytometry analysis of Tomato^+^ and Tomato^-^ hepatocyte populations for polyploidy. n=3. Data were presented as mean ±SD. Two-way ANOVA was applied. *P*-value <0.05 was considered as statistically significant.

We found that while 47.4% of Tomato^-^ hepatocytes were diploid cells, only 22.9% of Tomato^+^ hepatocytes were diploid in adult *Tomato^Twist2^* mice (Figure 3D), indicating that Twist2-lineage hepatocytes are largely polyploids. The percentage of polyploid hepatocytes was down from 77.1% to 68.9% in 20-month-old *Tomato^Twist2^* mice compared to 2-month old mice, although the percentage of Tomato^+^ hepatocytes in the liver did not significantly change with age (Figure S6), supporting that Twist2-lineage hepatocytes may undergo very slow turnover.

### Deletion of one *Twist2* allele did not affect midzone hepatocytes

To exclude the possibility that disruption of one allele of *Twist2* in *Twist2-Cre* mice affects hepatocytes in the midzone, we analyzed midzone hepatocytes using an established marker, Cyclin D1(Wei et al., 2021). We found that compared to age-matched wildtype mice, *Twist2-Cre* mice showed normal numbers of Cyclin D1-positive midzone hepatocytes at the age of P14 or P60 (Figure S7A, B). In addition, in CCl_4_-induced liver injury models, *Twist2-Cre* mice showed no alteration in Cyclin D1-positive midzone hepatocytes compared to wildtype mice (Figure S7C, D). These results suggest that the *Twist2-Cre* mouse line can be safely used to study midzone hepatocytes.

### Twist2-lineage hepatocytes show increased proliferation and migration capacities

We then directly compared Twist2-lineage hepatocytes (Tomato^+^) and conventional hepatocytes (Tomato^-^) from the same adult mouse by bulk RNA sequencing. KEGG pathway and GO term analyses revealed that Twist2-lineage hepatocytes were enriched in genes in mitogenic pathways, including Hippo, MAPK, Notch1, and TGFβ; ECM genes; and genes related to angiogenic regulation, cell migration, and cell shape and structure, but expression of metabolic genes, including *Alb* (encoding albumin), was greatly reduced (Figure 4A-D). Immunostaining confirmed increased activation of mTOR and Erks and increased expression of Notch downstream Hes1 in midzone hepatocytes (Figure 4E). Overall, these results suggest that Twist2-lineage hepatocytes might have greater mitogenic and migratory potentials at the sacrifice of metabolism. Indeed, we found that the percentage of proliferating Tomato^+^ hepatocytes were much higher than that in Tomato^-^ hepatocytes in P14 or adult mice (Figure 4F and later results).

**Fig. 4.**
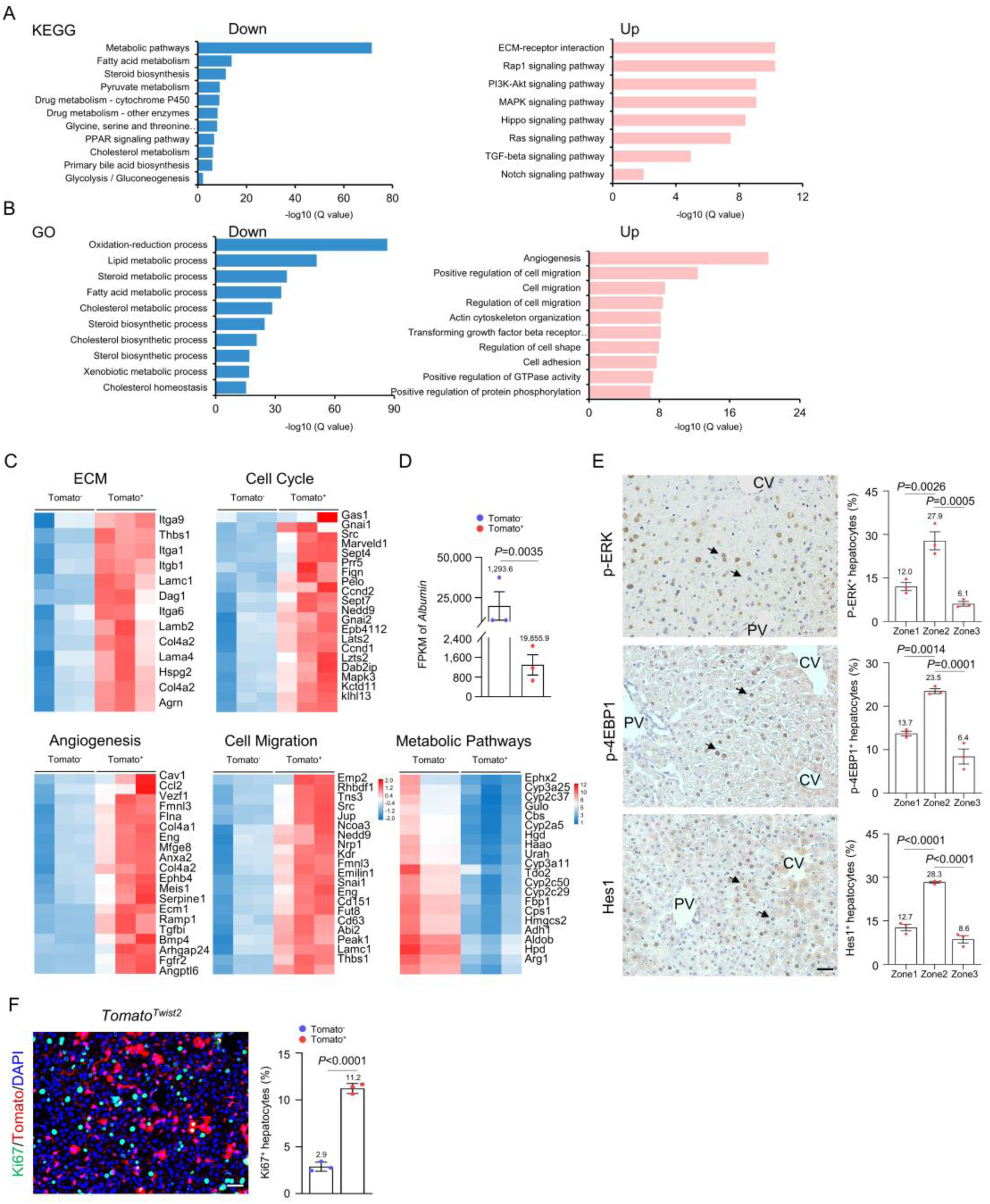
Transcriptome profiling of the mesenchymal-hepatocytes. **(A).** KEGG analysis of bulk RNA-seq data from Tomato^+^ and Tomato^-^ hepatocytes. **(B).** GO analysis of bulk RNA-seq data from Tomato^+^ and Tomato^-^ hepatocytes. **(C).** Heatmaps showing genes in the ECM, cell cycle, angiogenesis, cell migration, and metabolism modules of Tomato^+^ and Tomato^-^ hepatocytes. **(D).** FPKM values for *Alb* in Tomato^+^ and Tomato^-^ hepatocytes. n=3, Data were presented as mean ± SD. Unpaired two-tailed Student’s *t*-test was applied. *P*-value <0.05 was considered as statistically significant. **(E).** Analysis of p-ERK, p-4EBP1, and Hes1 on liver sections with immunohistochemical staining. Two-month-old normal mice were used. Arrows: pathway signals. Scale bar, 50 µm. Right panels; quantitation data. n=3. Data were presented as mean ± SD. Two-way ANOVA was applied. *P*-value <0.05 was considered as statistically significant. **(F).** Illustrative immunostaining results for Ki67^+^ cells among Tomato^+^ and Tomato^-^ hepatocytes of P14 mice. Hepatocytes were counted based on cell morphology and size. n=3. Scale bar, 50 µm. Data were presented as mean ±SD. Unpaired two-tailed Student’s *t*-test was applied. *P*-value <0.05 was considered as statistically significant.

### scRNA-seq analysis uncovers hepatocytes with mesenchymal features in adult mice

We then re-analyzed the public domain scRNA-seq datasets of the whole liver cells of adult mice based on *Alb* expression and found that hepatocytes could be divided into 4 subpopulations with 2 subgroups expressing about 5 times lower *Alb* than the other two (Figure S5A). Adult livers did not have the EpCAM^+^ populations observed in P1 and P14 mice (Figure S4A), suggesting that they are transit progenitors during liver growth. Alb^low^ subgroup 2 also expressed mesenchymal markers *Vimentin, Col3a1*, and *Col1a2* (Figure S4C, D). Moreover, comparative studies revealed that this Alb^low^ population showed enhanced expression of genes involved in liver development, cell cycle, growth, and features of Sertoli and adipocytes whereas the Alb^high^ groups expressed mainly ribosome, oxidation, ATP synthesis, and nucleotide metabolism (Figure S4E, F). These results suggest that the Alb^low^ 2 hepatocytes might overlap with the Twist2-lineage hepatocytes.

### Twist2-lineage hepatocytes show great transzone regeneration ability

To determine the possible roles of Twist2-lineage hepatocytes in liver regeneration, we first induced liver injury in *Tomato^Twist2^* mice with CCl_4_, which mainly damages zone 3 and causes fibrosis. In the acute phase, 2-3 times more Tomato^+^ hepatocytes were undergoing proliferation than Tomato^-^ hepatocytes (Figure 5A). When the injury had been repaired, the percentage of Tomato^+^ hepatocytes had increased to 39.1% from 12.0%, accompanied by conversion from polyploid to diploid hepatocytes (from 77.1% to 40.4%) (Figure 5B, C), consistent with previous studies showing the loss of polyploids in regenerated hepatocytes(Duncan et al., 2010). Conversion of polyploid to diploid hepatocytes may contribute to rapid expansion of Twist2-linegae hepatocytes during liver regeneration, since they do not need to duplicate the genome. Moreover, Tomato^+^ hepatocytes were detected close to the fibrotic Vimentin^+^ pericentral region (Figure 5B), indicating that they expanded to zone 3.

**Fig. 5.**
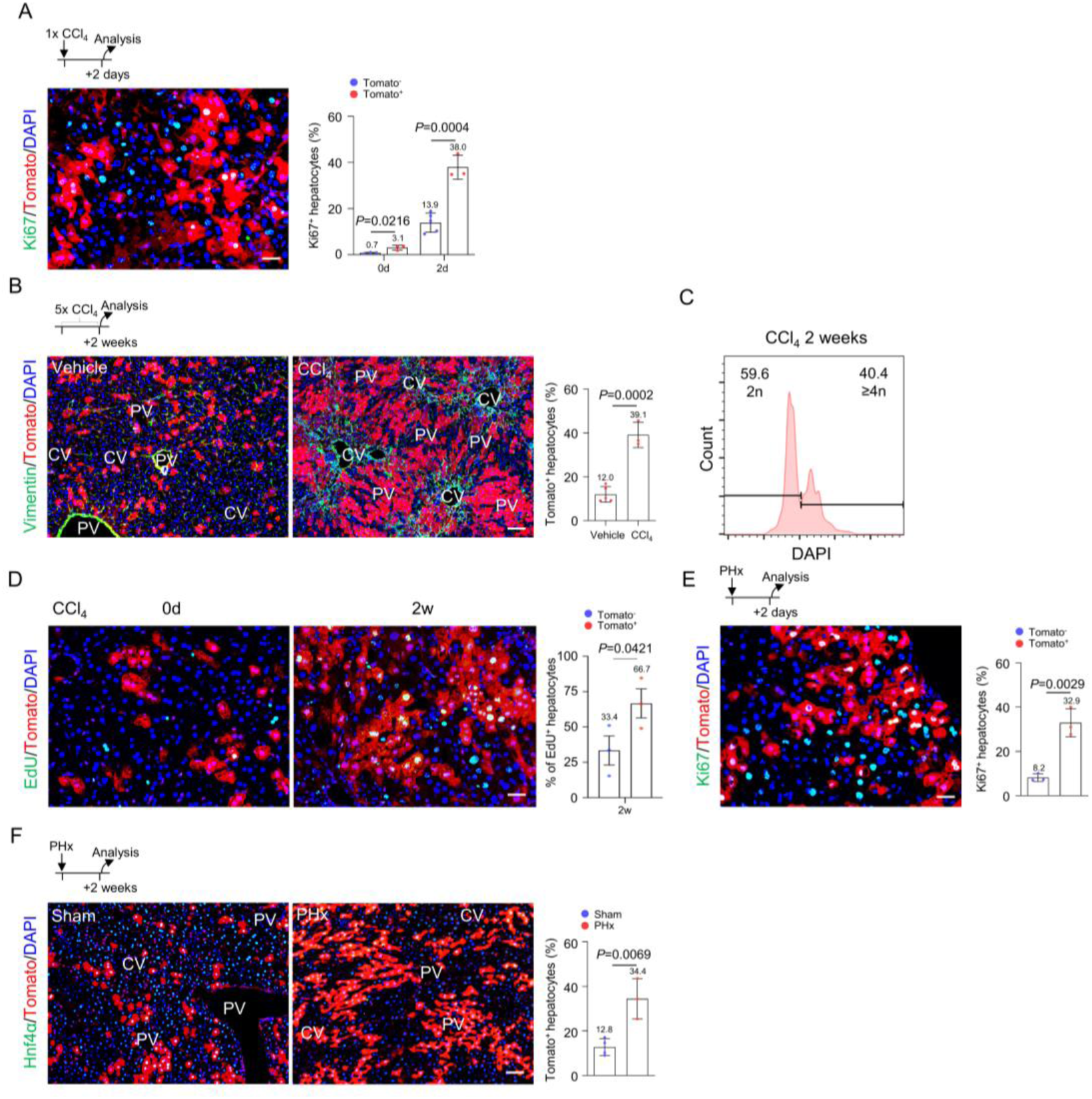
Mesenchymal-hepatocytes show greater regenerative ability than conventional hepatocytes. **(A).** Illustrative immunostaining results for Ki67^+^ cells among Tomato^+^ and Tomato^-^ hepatocytes of adult mice treated with CCl_4_ for 2 days. Right panels: quantitation data. n=5 for Tomato^-^ cells and n=3 for Tomato^+^ cells. Scale bar, 50 µm. Data were presented as mean ± SD. Two-way ANOVA was applied. *P*-value <0.05 was considered as statistically significant. **(B).** The percentage of Tomato^+^ hepatocytes in *Tomato^Twist2^* mice was increased 2 weeks after CCl_4_ administration. Vimentin was stained to detect fibrosis. Right panels: quantitation data. n=5 for Veh and n=3 for CCl_4_. Scale bar, 100 µm. Data were presented as mean ± SD. Unpaired two-tailed Student’s *t*-test was applied. *P*-value <0.05 was considered as statistically significant. **(C).** Changes in diploid and polyploid hepatocytes in regenerated livers of CCl_4_-induced injury model mice (*Tomato^Twist2^*). See Fig. 3d for control normal mice. **(D).** Immunostaining for the cumulative EdU-labelled cells among Tomato^+^ and Tomato^-^ hepatocytes of adult mice (14 days after CCl_4_), which received EdU every day to label all divided cells. Scale bar, 50 µm. Right panel: quantitation data. n=3. Data were presented as mean ±SD. Unpaired two-tailed Student’s *t*-test was applied. *P*-value <0.05 was considered as statistically significant. **(E).** Immunostaining results for Ki67^+^ cells among Tomato^+^ and Tomato^-^ hepatocytes from adult mice 2 days after PHx. Scale bar, 50 µm. Right panel: quantitation data. n=3. Data were presented as mean ± SD. Unpaired two-tailed Student’s *t*-test was applied. *P*-value <0.05 was considered as statistically significant. **(F).** The percentage of Tomato^+^ hepatocytes in *Tomato^Twist2^* mice was increased 2 weeks after PHx. Hnf4a was stained to show hepatocytes. Scale bar, 100 µm. Right panel: quantitation data. n=4 for sham and n=3 for PHx. Data were presented as mean ±SD. Unpaired two-tailed Student’s *t*-test was applied. *P*-value <0.05 was considered as statistically significant.

To validate the hyperproliferation of Twist2 lineage hepatocytes, we injected EdU to wildtype and *Tomato^Twist2^* mice daily for 14 days after CCl_4_-induced liver injury. We found that in EdU-labelled hepatocytes, 67% are Tomato^+^ (Figure 5D). These results confirmed that the Twist2 lineage mesenchymal-hepatocytes make greater contribution to liver regeneration than conventional hepatocytes.

We also tested the contribution of Twist2-lineage hepatocytes to liver regeneration in the PHx model, which mimics split liver transplantation in humans(Bangru & Kalsotra, 2020). We observed a 3-fold increase in proliferating Tomato^+^ hepatocytes at day 2 after 2/3 PHx and a 2 fold increase in the percentage of Tomato^+^ hepatocytes after regeneration, which also replenished zone 1 and, to a lesser extent, zone 3 (Figure 5E, F). Note that in the liver after 2/3 PHx, about 4% of Tomato^+^ hepatocytes remained, which generated 30.4% of new hepatocytes in the repaired liver, while 29% of Tomato^-^ hepatocytes remained, which generated 36.6% of new hepatocytes. Overall, these studies suggest that the midzone-located Twist2-lineage hepatocytes have greater proliferating and migrating capacities and regenerate about half of the new liver mass, while the conventional hepatocytes and stem/progenitor cells, which account for a vast majority of hepatocytes, regenerate the other half.

### Expression of *Twist2* or MET genes was unaltered during liver regeneration

To exclude the possibility that Twist2 expression is increased during liver regeneration, which may contribute to the increase in the number of Twist2-marked cells, we analyzed the public scRNA-seq data and found that the number of cells expressing Twist2 was not altered during liver regeneration, and Twist2 was mainly expressed in MCs rather than hepatocytes (Figure S8A, B). Immunostaining confirmed that Twist2 was mainly expressed in MCs rather than hepatocytes (with larger nuclei than MCs) (Figure S8B), suggesting that Twist2 expression is not altered during livre regeneration. We also tested whether MET (mesenchymal-epithelial transition) occurs during liver regeneration. We analyzed the expression of MET-related genes (*Snail*, *Twist1*, and *Zeb1*) and found that expression of these genes was not altered in regenerating livers of CCl_4_-induced injury or PHx models in wildtype or *Twist2-Cre* mice (Figure S8C, D). These results suggest that MET does not play a role in liver regeneration.

### Notch1 signaling suppresses mesenchymal-hepatocyte expansion and liver growth

Our RNA-seq data revealed that Twist2-lineage hepatocytes in adult mice showed enriched expression of 57 genes in the Notch1 pathway including *Hes1*, *Hey*, and *Heyl* (Figure 6A and Figure S9A). Notch molecules are required for liver development, especially cholangiocyte fate determination, and mutations in *Jag1* or *Notch2* cause a paucity of bile ducts in Alagille syndrome patients and mouse models(Dill et al., 2012; Fabris et al., 2019; Greenhill, 2014; Li et al., 1997; Romeo, 2019). We then generated *Notch1^Twist2^* mice (Figure S9B). These mutant mice appeared normal up to 12 months but had 4 instead of 7 liver lobes staring at birth, but showed an increased liver weight-to-body weight ratio (from 5.58% to 6.77%) (Figure 6B). The abundance of Tomato^+^ hepatocytes was increased by close to 2 folds, and these cells occupied zones 1 and 3 and showed enhanced proliferation (Figure 6C). These results suggest that Notch1 suppresses the proliferation of Twist2-lineage hepatocytes. It has been reported that Twist2 also marks BM-MSCs and muscle stem cells(Liu et al., 2017; Yu et al., 2003). However, *Nocth1* deletion did not affect skeletal structure, overall bone mass, or the weight or structure of skeletal muscles (Figure S9C, D), suggesting a tissue-specific effect for *Notch1* ablation on the liver.

**Fig. 6.**
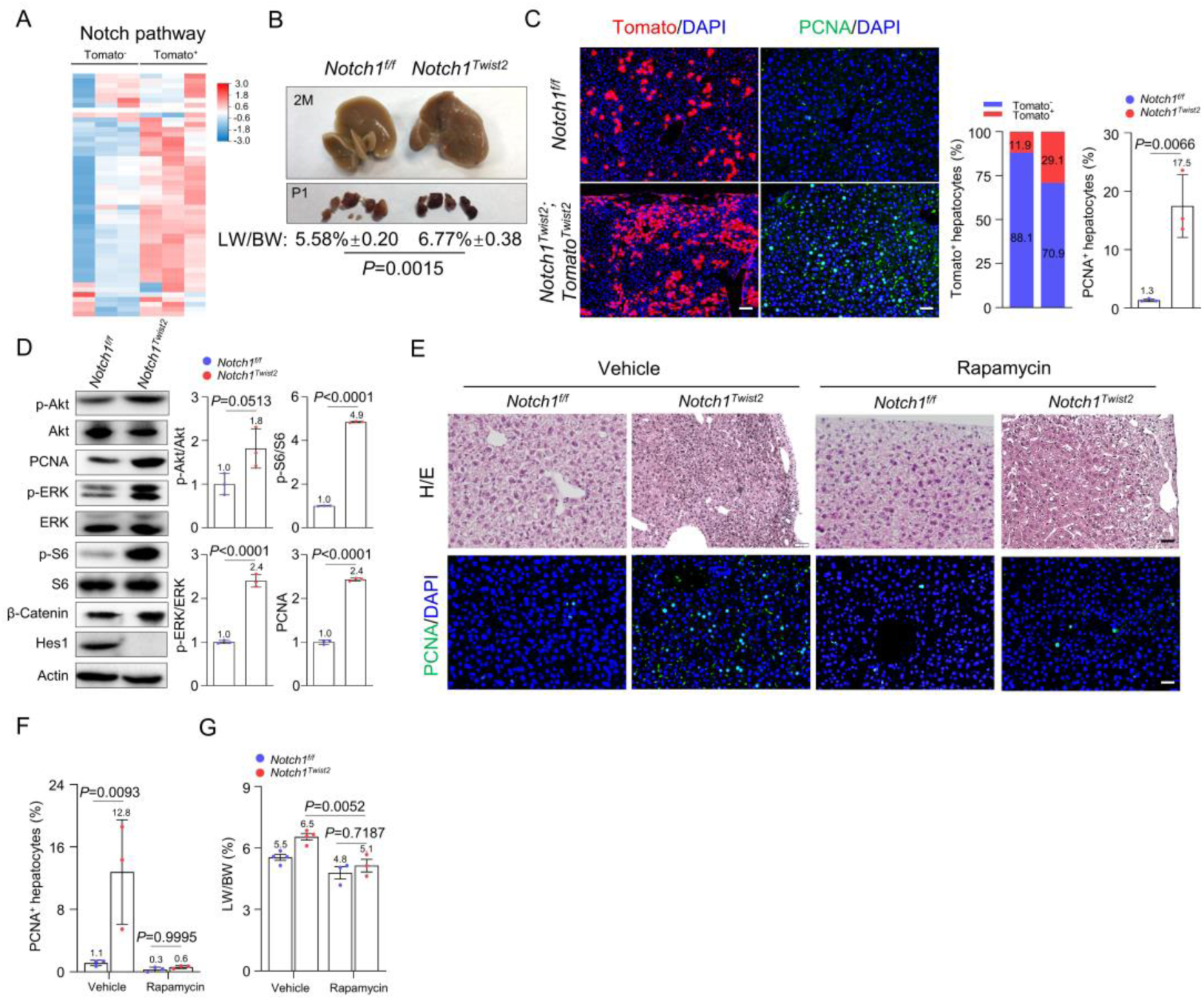
Notch1 suppresses mesenchymal-hepatocyte expansion via mTOR signaling. **(A).** RNA-seq data revealed that Twist2-lineage (Tomato^+^) hepatocytes are enriched in Notch1 pathway gene expression compared to Tomato^-^ hepatocytes. n=3. **(B).** *Notch1^Twist2^*mice had 4 instead of 7 liver lobes at birth and showed an increased liver weight-to-body weight ratio at adulthood. n=4. Data were presented as mean ± SD. Unpaired two-tailed Student’s *t*-test was applied. *P*-value <0.05 was considered as statistically significant. **(C).** Histological analysis revealed an increase in the number of Tomato^+^ hepatocytes and Ki67^+^ cells in adult *Notch1 ^Twist2^*;*Tomato^Twist2^* mice. Right panel: quantitation data. n=3. Scale bar, 100 µm. Data were presented as mean ± SD. Unpaired two-tailed Student’s *t*-test was applied. *P*-value <0.05 was considered as statistically significant. **(D).** Analysis of major mitogenic signaling pathways by Western blotting. Right panels: quantitation data. n=3. Data were presented as mean ± SD. Unpaired two-tailed Student’s *t*-test was applied. *P*-value <0.05 was considered as statistically significant. **(E).** RAP treatment diminished overproliferation in the livers of *Notch1^Twist2^* mice. n=3. Scale bar, 50 µm. **(F).** Quantitation data for proliferating cells. n=3. Data were presented as mean ±SD. Two-way ANOVA was applied. *P*-value <0.05 was considered as statistically significant. **(G).** RAP treatment restored the liver weight-to-body weight ratio to close-to-normal levels in adult *Notch1^Twist2^*mice. n=4 for Veh and n=3 for Rapamycin. Data were presented as mean ± SD. Two-way ANOVA was applied. *P*-value <0.05 was considered as statistically significant.

### Evidence that suppression of Notch1 signaling helps liver regeneration

Moreover, we found that during liver regeneration in normal mice, Notch1 signaling was suppressed in the midzone, manifested by a decrease in Hes1 signals (Figure S9E). Inhibiting of Notch signaling would promote proliferation of Twist2-lineage hepatocytes, as observed in *Notch1^Twist2^* mice (Figure 6C). We tested liver regeneration in *Notch1^Twist2^* mice and found that the percentage of Tomato^+^ hepatocytes did not increase further in CCl_4_-treated compared to non-treated mutant mice, this is in contrast to 2-3 fold increase in the percentage of Tomato^+^ hepatocyte in *Notch1^+/+^*mice (Figure S9F). Overall, these results suggest that suppression of Notch1 signaling in hepatocytes promotes expansion of the mesenchymal-hepatocytes during liver regeneration.

### Inhibiting VEGFR or mTOR diminishes *Notch1* ablation-induced hepatocyte expansion

Western blot analysis showed liver overgrowth in *Notch1^Twist2^*mice was associated with increased activation of the mitogenic signaling molecules Akt1, Erk, and mTOR in Notch1-deficient liver samples (Figure 6D). Immunostaining confirmed strong activation of ERKs and mTOR on liver sections (Figure S9G). mTOR is a sensor of nutrients and growth factors and can be activated by both Akt1 and ERK(Lu et al., 2021). We found that treatment of *Notch1^Twist2^* mice with rapamycin (RAP), an mTOR complex 1 inhibitor, for 1 month starting at 1 month of age restored the liver structure, the percentage of PCNA^+^ hepatocytes, and the LW/BW ratio to close-to-normal levels, yet, RAP did not significantly affect the liver structure or LW/BW ratio in normal mice (Figure 6E-G), suggesting that enhanced mTOR signaling contributes to hyperproliferation of Notch1-deficient mesenchymal-hepatocytes.

The livers of *Notch1^Twist2^* mice also showed increased angiogenesis (Figure S9H). Previous studies have shown that hepatocytes secrete VEGF to promote angiogenesis(Banerjee et al., 2015; Cuervo et al., 2016; Dill et al., 2012), as well as hepatocyte proliferation(LeCouter et al., 2003). We found that *Notch1-*deficient liver samples showed enriched expression of pro-angiogenesis factors, especially *Vegfa* and *Vegfb* (Figure 4C and 7A). Previous studies have shown that Notch1 inhibits *Vegf* expression and angiogenesis, likely via Hif1α (Θιανγ ετ αλ., 2012; Zηυ ετ αλ., 2017). Our immunostaining experiments uncovered an increase in Hif1α in hepatocytes of *Notch1^Twist2^* mice (Figure 7B), which was mainly nuclear, supporting that *Notch1* signaling suppresses *Vegfa* expression. Functionally, treating *Notch1^Twist2^* mice with axitinib, an inhibitor of VEGF receptors, for 1 month starting at the age of 1 month diminished the increases in angiogenesis, the number of PCNA^+^ proliferating hepatocytes, and the LW/BW ratio, as well as mTOR activation (Figure 7C-E). We also found that axitinib could inhibit proliferation of Huh7 cells, a hepatoblastoma cell line, in vitro (Figure S10). The findings that axitinib showed insignificant effect on the liver parameters in normal mice (Figure 7C-E), suggest that VEGF promotes hepatocyte proliferation when expressed at high levels.

**Fig. 7.**
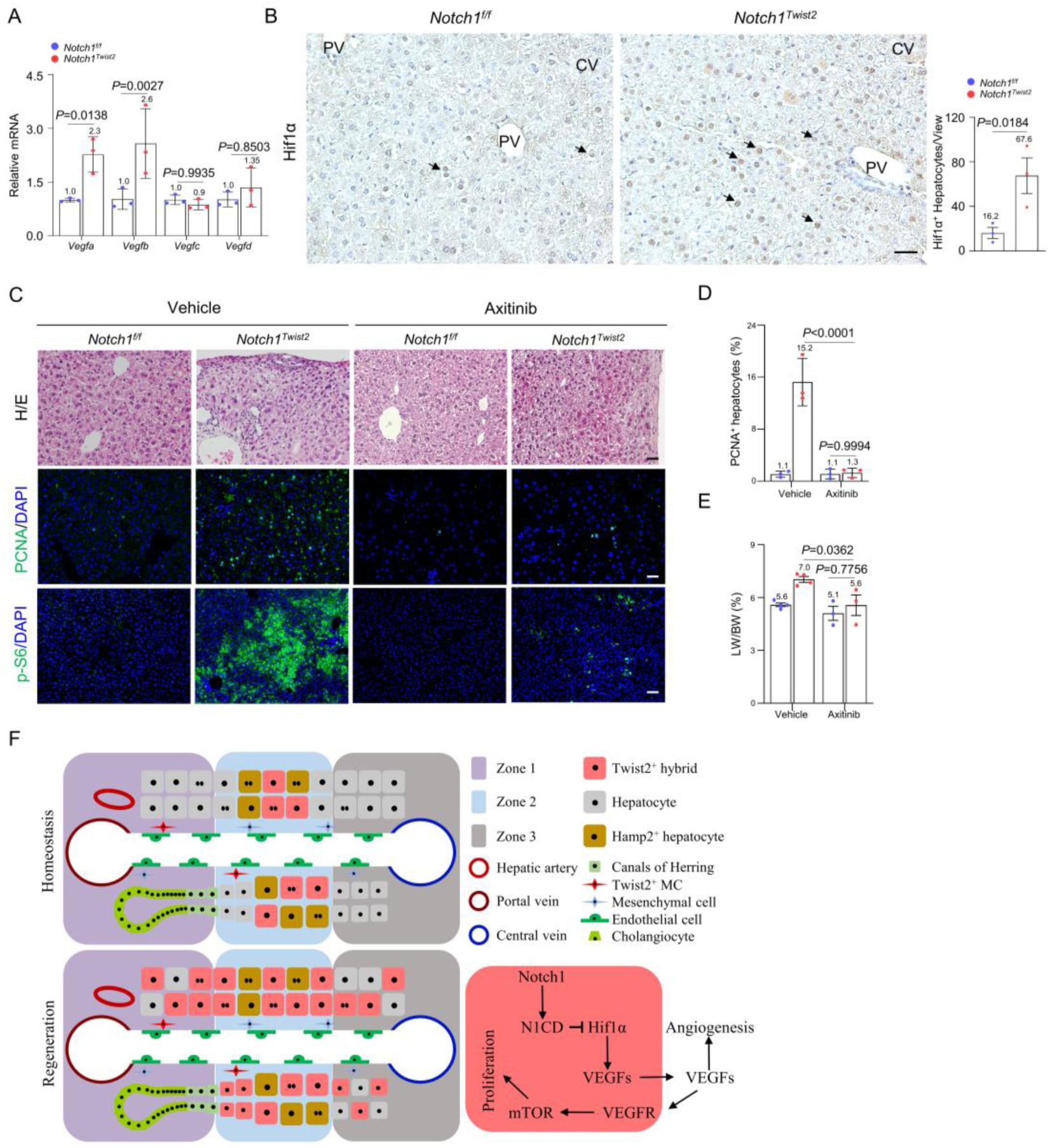
VEGFR inhibitor suppresses mTOR signaling and Twist2-lineage hepatocyte expansion in *Notch1^Twist2^* mice. **(A).** Liver samples from *Notch1^Twist2^*mice showed an increase in *Vegfa* and *Vegfb* expression but not *Vegfc* or *Vegfd* expression. The values of control samples were set at 1.0. n=3. Data were presented as mean ± SD. Two-way ANOVA was applied. *P*-value <0.05 was considered as statistically significant. **(B).** Immunostaining of Hif1α on liver sections. Arrows: Hif1α signals. Scale bar, 50 µm. Arrows indicate Hif1α signals. Right panel: quantitation data. n=3. Data were presented as mean ± SD. Unpaired two-tailed Student’s *t*-test was applied. *P*-value <0.05 was considered as statistically significant. **(C).** Axitinib treatment diminished overproliferation and mTOR activation in the livers of *Notch1^Twist2^*mice. n=3. Scale bar, 50 µm, 100 µm for p-S6. **(D).** Quantitation data for proliferating cells in livers among vehicle and axitinib groups. n=3. Data were presented as mean ± SD. Two-way ANOVA was applied. *P*-value <0.05 was considered as statistically significant. **(E).** Axitinib treatment restored the liver weight-to-body weight ratio to close-to-normal levels. n=4 for Veh and n=3 for Axitinib group. Data were presented as mean ± SD. Two-way ANOVA was applied. *P*-value <0.05 was considered as statistically significant. **(F).** A model depicting how Twist2 lineage mesenchymal-hepatocytes regenerate the liver in mice.

## Discussion

The liver is the only organ that can be fully regenerated in mouse and human. Rapid recovery 2/3 of the mouse liver mass requires not only massive expansion of remaining hepatocytes but also cell migration and structure rebuilding. Although adult stem cells play critical roles in regeneration of many tissues including the gut, skin, and skeletal muscles, cumulative evidence suggests that the differentiated hepatocytes, especially midzone hepatocytes, play a dominant role in liver regeneration(Andersson, 2021; Monga, 2020). However, these cells are terminally differentiated epithelial cells and in general have limited proliferation and migratory potentials. So how liver is rapidly regenerated remains not fully resolved.

Our current study uncovered a small mesenchymal-hepatocyte hybrid population, which possess the features required for rapid liver regeneration. These cells are likely derived from Twist2-expressing cells in embryonic livers via the intermediate EpCAM^+^ cells in p7 and P14 pups(Lotto et al., 2020). scRNA-seq analysis suggest that such hepatocytes exist in adult mice. Certainly, the ontology of this lineage warrants further investigation. The Twist2 lineage hepatocytes express higher levels of mesenchymal-related genes and lower levels of metabolic genes compared to conventional hepatocytes, are mainly located in the midzone, highly polyploidy, and display increased proliferation and migration capacities. A recent study also uncover these features for midzone hepatocytes(Sun et al., 2020). During liver regeneration, these cells showed 2-3 times greater proliferation capacity than conventional hepatocytes and the percentages of these cells almost tripled after repair. Moreover, they replenished not only the midzone but also zones 1 and 3. Overall, these findings suggest that the mesenchymal-hepatocytes represent an important regenerative force and form about half of the new liver mass (Figure 7F).

The mesenchymal-hepatocytes appear to play a role in liver growth but not much in liver homeostasis. During postnatal liver growth, these cells undergo proliferation at a rate greater than that of conventional hepatocytes. Moreover, deletion of *Notch1* in these cells led to expansion of the mesenchymal-hepatocytes by 3 folds, which repopulate zone 1 and 3, accompanied by an increase in the LW/BW ratio, suggesting that these cells participate in liver postnatal growth, which is under the control of Notch1 signaling. However, in adult mice, the size of the mesenchymal-hepatocyte population did not change up to 20 months of age. Despite that, we observed a modest loss of polyploid hepatocytes, suggesting that they may undergo slow turnover.

An intriguing finding is that most of the highly mitogenic and migratory mesenchymal-hepatocytes are polyploidy and moreover, they show a conversion from polyploid to diploid with age or during regeneration, which may contribute to rapid expansion of Twist2-lineage hepatocytes as polyploid cells do not need to duplicate the genome before division, consistent with previous studies showing that polyploid hepatocytes turn into diploids during regeneration and they undergo little DNA synthesis(Duncan et al., 2010; Wilkinson et al., 2019). Certainly, the function of diploid conversion warrants further investigation.

Although our findings confirm the important role played by midzone hepatocytes in liver regeneration, they also suggest that not all hepatocytes are equal in the midzone. Hamp2 is specifically expressed in some midzone hepatocyte. Twist2-lineage hepatocytes do not express Hamp2 and differ from Hamp2^+^ hepatocytes, as the latter undergoes constant turnover in adult mice and expands much less during regeneration (from 10% to 13% in the CCl_4_ injury model and from 10 to 17.5% in the diethoxycarbonyl-1,4-dihydrocollidine-induced model)(Wei et al., 2021). Moreover, our study of the PHx model suggests that conventional hepatocytes also participate in liver regeneration, although at a slow pace, consistent with previous findings(Monga, 2020). Cooperation of mesenchymal-hepatocytes and conventional hepatocytes are thus needed for rapid liver regeneration.

We also show that Notch1 activation is increased in Twist2-lineage hepatocytes compared to conventional hepatocytes and that this activation is suppressed during liver regeneration. Deletion of *Nocth1* led to the expansion of these hepatocytes into zones 1 and 3 and an increase in the liver weight-to-body weight ratio. This is in contrast to mice in which *Nocth1* was ablated from Alb^+^ hepatocytes, which did not develop obvious liver phenotypes(Geisler et al., 2008). This can be explained by late and weak *Alb* expression in Twist2-lineage hepatocytes, restriction of the function of Notch1 to progenitor cells, and/or weak Notch1 activation in conventional hepatocytes (Figure 6A). Our pharmacological studies suggest that Nocth1 may negatively regulates the expression of *Vegfa* and *Vegfb* in Twist2-lineage cells, which promote cell proliferation via VEGFR-mTOR signaling in autocrine and/or paracrine manners.

One complication in using knock-in Cre or CreERT mice is that ablation one allele of the marker gene may affect the behavior of the labelled cells. Here we show that ablation one Twist2 allele did not affect liver development, growth, or regeneration, nor does it affect the midzone hepatocytes. Moreover, we show that expression of Twist2 or MET related genes is not affected during liver regeneration, and that deletion of one *Twist2* allele does not affect the expression of MET genes. These results indicate that it is safe to use the *Twist2-Cre* to study liver development and regeneration.

In summary, the findings of this study suggest that in response to injury-induced cues, midzone-located mesenchymal-hepatocytes undergo great expansion and migration and regenerate zones 1 and 3, which form half of the new liver mass. Moreover, the highly regenerative and mesenchymal nature of this hepatocyte subpopulation implies that they may constitute a cell basis for liver cancer, fat liver, and liver cirrhosis, thus opening up a new direction in the study of liver diseases.

## Materials and Methods

### Mice

The *Twist2-Cre* (008712), *Notch1^f/f^* (006951)*, R26-tdTomato* (007914), *Col1α2-CreERT* (029567), and R26-*mTmG* (007576) mouse lines were obtained from the Jackson Laboratory. The *Acta2-Cre* line was generated in Xiao Yang’s laboratory. The knock-in *Twist2^f/f^*and *Vim-CreERT* line were generated by Shanghai Biomodel Organism Science & Technology Development Co., Ltd. The *CreERT* cassettes was inserted in front of the ATG codon. DNA sequencing confirmed proper recombination at the locus. All mice were housed in the animal facility at Shanghai Jiao Tong University, and mouse work was performed following the recommendations of the National Research Council Guide for the Care and Use of Laboratory Animals with a protocol approved by the Institutional Animal Care and Use Committee of Shanghai, China [SYXK (SH) 2011-0112]. For tracing and gene deletion experiments, two-month-old male mice were used.

### Histology, immunohistochemistry, and immunofluorescence

Mice were perfused with 4% paraformaldehyde (PFA), and the organs were harvested immediately after sacrifice, washed, embedded in paraffin, sectioned at 5 μm, and stained with hematoxylin and eosin (H/E). For immunostaining, antigen retrieval was performed by boiling the slides in citrate buffer (pH 6) for 10 min, followed by cooling to room temperature. Tissues were submerged in methanol containing 3% H_2_O_2_ for 20 min to inactivate endogenous peroxidases, followed by treatment with 0.1% Triton X-100 for 20 min to permeabilize the cells. The sections were blocked in 10% goat serum for 45 min and incubated with primary antibodies at 4°C overnight and secondary antibodies (Invitrogen) for 1 h at 37°C.

To prepare frozen sections, tissues were collected in cold PBS and then fixed in 4% PFA at 4°C for 4 hours. After washing in PBS three times, the tissues were placed in 30% sucrose/PBS overnight at 4°C, embedded in OCT compound, and stored at -80°C until sectioning. Cryosections (10 µm in thickness) were collected on positively charged slides and stored at -20°C until use. The following antibodies were used: p-S6 (1:100, CST, 2211), PCNA (1:100, CST, 2586), p-ERK (1:100, CST, 9106), Ki67 (1:100, Thermo Scientific, PA5-19462), p-4EBP1 (1:100, CST, 2855), Hif1α (1:100, Invitrogen, PA1-16601), Hes1 (1:100, Abcam, 71559). PDGFRα (1:100, R&D, AF1062), CD31 (1:50, Abcam, 7388), CD45 (1:100, Abcam, 154885), Hnf4α (1:100, Abcam, 181604), K19 (1:100, DSHB, Troma-III), αSMA (1:100, Abcam,119952), Vimentin (1:100, Abcam, 193555), Cdh1 (1:100, R&D, AF748), Cyp2e1 (1:100, Sigma, HPA009128), Col1 (1:100, Abcam, 260043). Twist2 (1:50, Sigma, WH0117581M1), Cyclin D1 (1:100, Abcam, 16663).

### Liver injury model

For the CCl_4_-induced liver injury model, mice were intraperitoneally injected with CCl_4_ (diluted 1:4 in corn oil) at a final volume of 5 ml/kg body weight. For the PHx liver injury model, 2/3 PHx injury was performed according to established protocols. Briefly, mice were anesthetized by an intraperitoneal injection of 40 mg/kg sodium pentobarbital and transferred onto an electric blanket. Following removal of the abdominal fur and skin disinfection with iodine, a midline abdominal skin and muscle incision was made to expose the liver. The base of the left lateral lobe and the median lobe were ligated with a 4-0 silk thread, and the lobes were resected just above the knots. The peritoneum and skin were then closed with 6-0 sutures, and the mice were supplied with pure oxygen for 4-5 minutes until they achieved normal breathing. Then, the mice were placed in individual cages under a warming lamp for recovery. The right and caudate lobes were used for analysis.

### X-ray imaging of the skeleton

Bone radiographs were taken with a cabinet X-ray system (LX-60, Faxitron Bioptics) using standardized settings (45 kV for 8 s).

### Determination of long term hepatocyte proliferation

EdU (5-Ethynyl-2-deoxyuridine) (5 mg/kg) was injected intraperitoneally into *Tomato^Twist2^*mice daily for 14 days after CCl_4_-induced liver injury. Cell-LightTM Apollo488 Stain Kit(100T) (Code No. C10371-3) (RIBOBIO, China) was used to detect EdU incorporation into the cellular DNA. Briefly, after embedding, the frozen sections were washed in phosphate-buffered saline for 15 minutes at room temperature and incubated with 2 mg/ml glycine buffer for 10 minutes at room temperature. After washing with 0.5% triton x-100 buffer for 10 minutes, a mixed reaction buffer was added to samples (30 min at room temperature). Staining of the nuclei was performed with DAPI.

### RNA isolation and quantitative PCR analysis

Total RNA was isolated using TRIzol (Invitrogen), and 1 μg of total RNA was used to synthesize complementary DNA using random primers (SuperScript II RT; Invitrogen). For quantitative PCR, SYBR Green PCR Master Mix (Applied Biosystems) was used in a CFX96 real-time thermocycler system (Bio-Rad); the RNA levels of target genes were normalized to those of GAPDH. For each qPCR experiment, all samples were run in triplicate. The following primers have been used in other studies:

*GAPDH*-F: 5’-TGACCTCAACTACATGGTCTACA-3’

*GAPDH*-R: 5’-CTTCCCATTCTCGGCCTTG-3’

*Vegfb*-F: 5’- GCCAGACAGGGTTGCCATAC -3’

*Vegfb*-R: 5’- GGAGTGGGATGGATGATGTCAG -3’

*Vegfc*-F: 5’- GAGGTCAAGGCTTTTGAAGGC -3’

*Vegfc*-R: 5’- CTGTCCTGGTATTGAGGGTGG -3’

*Vegfa*-F: 5’- GCACATAGAGAGAATGAGCTTCC -3’

*Vegfa*-R: 5’- CTCCGCTCTGAACAAGGCT -3’

*Vegfd*-F: 5’- TTGAGCGATCATCCCGGTC -3’

*Vegfd*-R: 5’- GCGTGAGTCCATACTGGCAAG -3’

*Hes1*-F: 5’- CATTCCAAGCTAGAGAAGGCAG-3’

*Hes1*-R: 5’- TATTTCCCCAACACGCTCG-3’

*Notch1*-F: 5’- TGCCAGGACCGTGACAACTC-3’

*Notch1*-R: 5’- CACAGGCACATTCGTAGCCATC-3’

*Snai1-*F: 5’- CACACGCTGCCTTGTGTCT-3’

*Snai1-*R: 5’-GGTCAGCAAAAGCACGGTT-3’

*Zeb1-*F: 5’-GCTGGCAAGACAACGTGAAAG-3’

*Zeb1-*R: 5’-GCCTCAGGATAAATGACGGC-3’

*Twist1-*F: 5’-GGACAAGCTGAGCAAGATTCA-3’

*Twist1-*R: 5’-CGGAGAAGGCGTAGCTGAG-3’

### Western blot analysis

All mouse liver tissues were ground and lysed in protein lysis buffer for use in Western blotting. The following antibodies were used: p-S6 (1:1000, CST#2211), S6 (1:1000, CST, 9211), Akt (1:1000, CST, 9272), p-Akt (1:1000, CST, 4060), PCNA (1:1000, CST, 2586), p-ERK (1:1000, CST, 9106), ERK (1:1000, CST, 9102), β-Catenin (1:1000, Santa Cruz Biotechnology, 7199), Hes1 (1:1000, Abcam, 71559), Actin (1:5000, Santa Cruz Biotechnology, 81178).

### Drug treatments

RAP (Selleck) was formulated in 0.25% PEG400, 0.25% Tween 80 and 99.75% water; this formulation was intraperitoneally injected at a dosage of 10 ml/kg daily for 30 days. Axitinib (Selleck) was dissolved in DMSO and formulated in 44.44% PEG300, 5.56% Tween 80 and 44.44% water; this formulation was administered by intraperitoneal injection at 7.5 ml/kg daily for 30 days.

### Liver cell isolation

Primary mouse cells (including parenchymal and mesenchymal cells) were isolated by the two-step liver perfusion method with modifications. Briefly, mice were anesthetized by an intraperitoneal injection of 40 mg/kg sodium pentobarbital. The abdomen was then cut open to expose the liver, inferior vena cava and portal vein. The portal vein was cut, and the liver was first perfused with 1x PBS (containing 0.5 mM EDTA) for 5 minutes and then perfused with digestion buffer containing collagenase type IV (Sigma) and DNase I for 7-10 minutes. The livers were then clipped, removed and transferred to a Petri dish filled with 1x PBS at 4°C. The liver tissues were gently torn to help the hepatocytes dissociate, and the cell suspension was filtered through a 70-μm cell filter (BD Falcon), followed by centrifugation at 4°C and 50 x g and 500 x g for hepatocytes and nonparenchymal cells, respectively. We then resuspended the cells in 50% and 25% Percoll mixtures and centrifuged them at 300 x g for 20 min at 4°C. Purified hepatocytes and nonparenchymal cells were collected for ploidy analysis or washed with PBS and resuspended in DMEM containing 10% FBS for cell culture.

### CCK-8 assays

Axitinib was added to the cells and cultured for different periods of time. Each well was then added with 10 μl CCK-8 reagent (Keygen Biotech, CN) and cultured for 3 more hrs. The viable cells were determined by measuring the optical density (OD) absorbance at the wavelength of 450 nm.

### Bulk RNA sequencing analysis

DNase I was used to digest double-stranded and single-stranded DNA in total RNA, and magnetic beads were then used to recover the reaction products. RNase H or the Ribo-Zero method (human, mouse, plants) (Illumina, USA) was used to remove rRNA. Purified mRNA from previous steps was fragmented into small pieces with fragment buffer at the appropriate temperature. Then, first-strand cDNA was generated in the First Strand Reaction System by PCR, and second-strand cDNA was generated as well. The reaction product was purified by magnetic beads, after which A-tailing mix and RNA index adapters were added to carry out end repair. The cDNA fragments with adapters were amplified by PCR, and the products were purified with Ampure XP beads. The library was validated on an Agilent Technologies 2100 bioanalyzer for quality control. The double-stranded PCR products were then heat-denatured and circularized by the splint oligo sequence. Single stranded circular DNA (ssCir DNA) was formatted as the final library. The final library was amplified with phi29 (Thermo Fisher Scientific, MA, USA) to prepare DNA nanoballs (DNBs) containing more than 300 copies of one molecule. The DNBs were loaded into the patterned nanoarray, and single-end 50-base reads were generated on the BGISEQ-500 platform (BGI-Shenzhen, China).

Differentially expressed genes (DEGs) determined from the RNA expression data were mapped to the GO database or KEGG database, and significant GO and KEGG pathway enrichment analyses were performed using the R package clusterProfiler.

### Flow cytometry and fluorescence-activated cell sorting (FACS)

For analysis of Tomato^+^ cells, Wild-type mouse cells were used to set compensation. DNA content was quantified with DAPI (Thermo Scientific)(Matsumoto et al., 2020). All flow cytometry analyses were performed on a Beckman CytoFLEX S. All flow cytometry sorting was performed on a Bio-Rad S3e. The FACS data were analyzed with FlowJo.

### 10X Illumina single-cell RNA sequencing

Digested tdTomato^+^ total liver cells from P1 and P14 *tdTomato^Twist2^* mice were sorted and used for sequencing. The cell suspension was loaded into Chromium microfluidic chips with 3’v3 chemistry and barcoded with a 10X chromium controller (10X Genomics). RNA from the barcoded cells was subsequently reverse-transcribed, and sequencing libraries were constructed with reagents from a Chromium Single-Cell 3’v3 Reagent Kit (10X Genomics) following the manufacturer’s instructions. Sequencing was performed with an Illumina NovaSeq 6000 (Illumina).

### Single-cell RNA analysis

Raw reads were demultiplexed and mapped to the mouse reference genome with the Cell Ranger version 3.0.1 (10X Genomics) pipeline using the default parameters. The generated gene-cell expression matrix was used for subsequent analysis in R version 3.6.1 using Seurat version 3.1.5(Butler et al., 2018). “Cells” with any of the following criteria were excluded: <500 expressed genes; >25% unique molecular identifiers (UMIs) mapped to mitochondria; or nUMI >99% events, with the number of UMIs (nUMI) used to exclude doublet cells. Clusters with very few cells were filtered before downstream analysis. Filtered cells from P1 and P14 samples were merged using the “merge” function in Seurat, and the merged data were used for downstream graph-based clustering and t-SNE visualization. DEGs in each cluster were identified with the function “FindAllMarkers” in Seurat. The “DoHeatmap” function in Seurat was used to obtain heatmap figures. The “FeaturePlot” and “VlnPlot” functions in Seurat were used to visualize specific gene expression.

For correlation analysis, average gene expression data for each cluster were generated from merged data. Correlation indexes between each cluster were calculated based on the average gene expression data using “cor” in R and drawn by the ggplot2 R package.

Using the marker genes listed in fig. S4, we calculated ‘scores’—a single numeric value representative of the expression of multiple marker genes—as the sum of log2(counts+1) across all markers in a category.

scRNA-seq data of MCs (GSE137720) and hepatocytes (GSE125688) in the liver were obtained from the GEO database. The analytical strategy was similar to that mentioned above. For DEGs between Alb^low^ 2 and Alb^high^ 1 & 2 cells from healthy hepatocyte samples, “FindMarkers” in Seurat was used.

### GO and KEGG pathway enrichment analyses

DEGs were mapped to the GO and KEGG databases, and enrichment analyses of GO terms and KEGG pathways were performed using the clusterProfiler R package(Yu et al., 2012).

### Pseudotime analysis

Merged Seurat data without “proerythroblasts” were used for pseudotemporal analysis using the Monocle2 R package(Qiu et al., 2017). The “differentialGeneTest” function was used to identify DEGs between the cells belonging to P1 samples and P14 samples. Ordering genes were selected from among DEGs with a q value cutoff of <1 × 10−100, which produced a list of 2562 genes. The “reduceDimension” function was used with the parameter “max_components = 2, method = ’DDRTree’”. “orderCells” was then run, and the result was graphed with the function “plot_cell_trajectory”. The branch containing the most P1 cells was set as the original root.

### RNA-seq data in public databases

The following RNA-seq datasets (available in public databases) were used in this study. Hepatocytes (GEO database accession: GSE125688) and mesenchymal cells (GEO database accession: GSE137720).

### Data available

scRNA-seq data have been deposited in the SRA database under accession code PRJNA721304. Bulk RNA-seq data of Twist2-linegae hepatocytes and conventional hepatocytes have been deposited in the SRA database under accession code PRJNA721538.

### Statistics

Data are represented as means ± S.D. The number of mice used for each experiment (without randomization) is indicated in the corresponding figure legend. Analyses of significant differences between groups were performed using two-tailed Student’s *t*-tests or Two-way ANOVA.

## Acknowledgement

The work was supported by the National Key Research and Development Program of China (2018YFA0800803 to BL and 2018YFA0800803 to JL) and the National Natural Science Foundation of China (81520108012 and 91749201) to BL.

## Author contributions

B. L. and J. L. designed research; G. Y., S. Z., A. R., and S. B. performed research; H. L and H. W. analyzed data; J. L. and B. L. wrote the paper.

## Supplementary Materials for

### Supplementary Figures

**Fig. S1.**
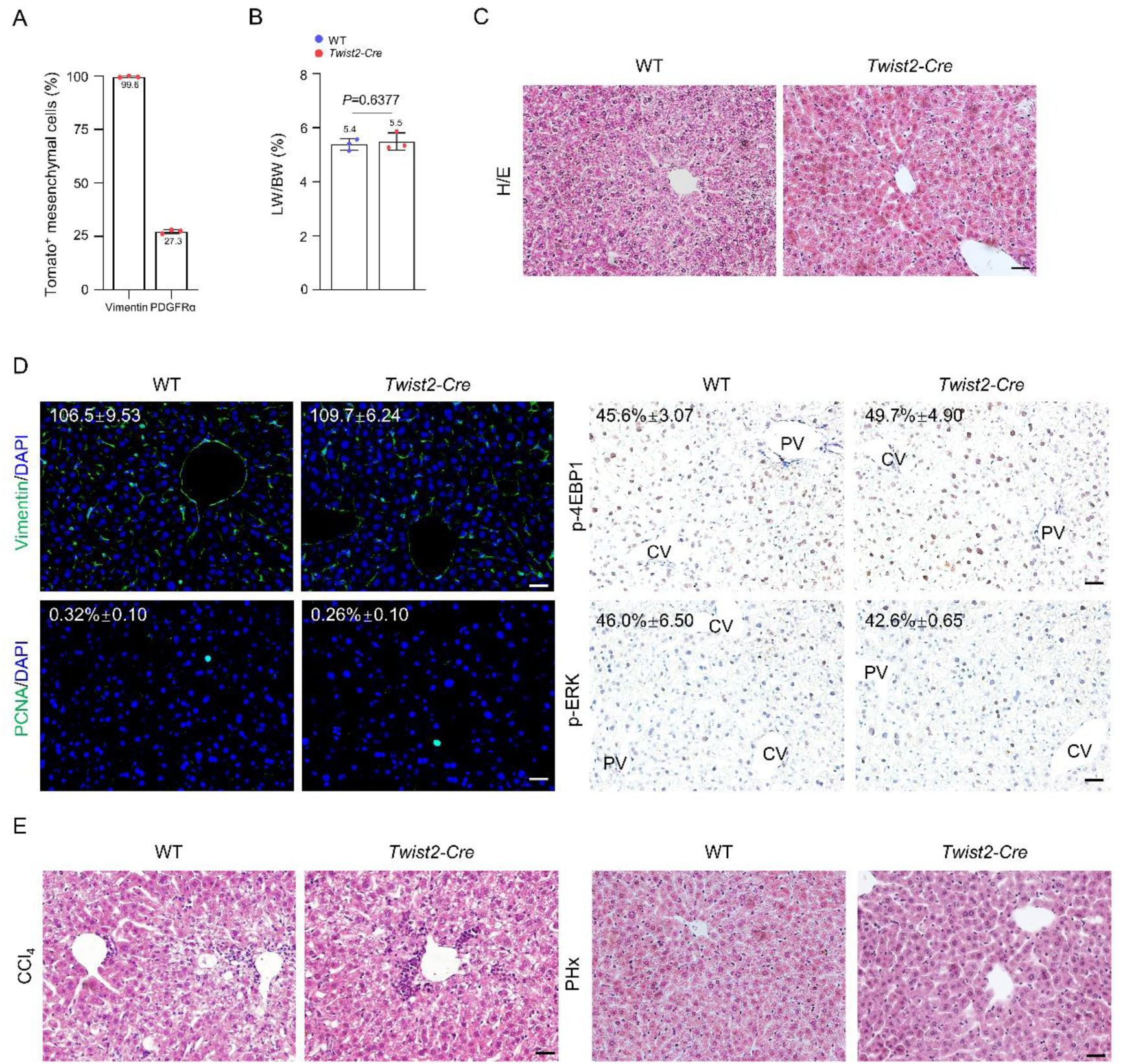
Analysis of liver development and regeneration in *Twist2-Cre* mice. **(A).** The percentages of Tomato^+^ MCs positive for Vimentin or PDGFRα (See immunostaining results in Fig. 1b). n=3 mice. **(B).** The liver weight-to-body weight ratios of adult male *Twist2-Cre* and control mice. n=3. Data were presented as mean ±SD. Unpaired two-tailed Student’s *t*-test was applied. *P*-value <0.05 was considered as statistically significant. **(C).** Histological structures of the livers of adult *Twist2-Cre* and control mice. Scale bar, 50 µm. **(D).** Immunostaining results for Vimentin, PCNA, p-4EBP1 and p-ERK on liver sections of adult *Twist2-Cre* and control mice. Quantitation data: the numbers of Vimentin^+^ cells per view and the percentages of cells positive for PCNA, p-4EBP1, or p-ERK. n=3. Scale bar, 50 µm. Data were shown as mean ±SD. **(E).** Illustrative H/E staining results of regenerated livers from *Twist2-Cre* and control mice 2 weeks after CCl_4_-induced injury or PHx. n=3. Scale bar, 50 µm.

**Fig. S2.**
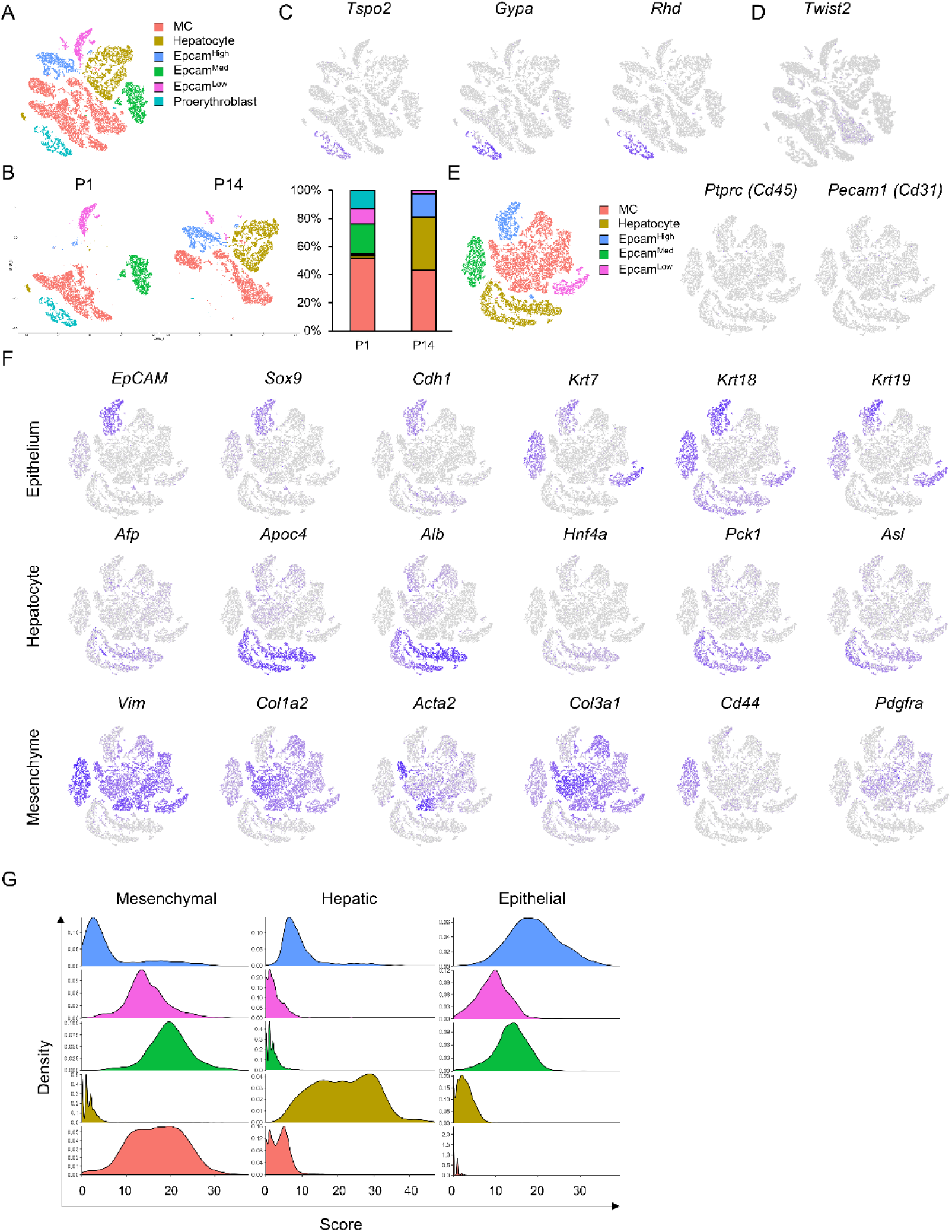
scRNA-seq analysis of liver Tomato^+^ cells from P1 and P14 mice. **(A).** t-SNE analysis of Tomato^+^ liver cells including proerythrocytes (combined Tomato^+^ cells from P1 and P14 *Tomato^Twist2^* mice). **(B).** Comparison of the Tomato^+^ subgroups from P1 and P14 mice. Right panels: percentage of each subpopulation. **(C).** t-SNE analysis showing the expression of proerythrocyte marker genes. **(D).** t-SNE analysis showing the expression of *Twist2* in various subpopulation. **(E).** t-SNE analysis of CD31 and CD45 expression in various Twist2 lineage groups exclusive of proerythrocytes. **(F).** t-SNE analysis of expression of important cell-specific marker genes for MCs, hepatocytes, and epithelial cells. **(G).** Comparison of the expression of multiple marker genes in different Tomato^+^ subgroups. Density indicates the percentage of cells with a given score and score indicates the equivalence to a given lineage.

**Fig. S3.**
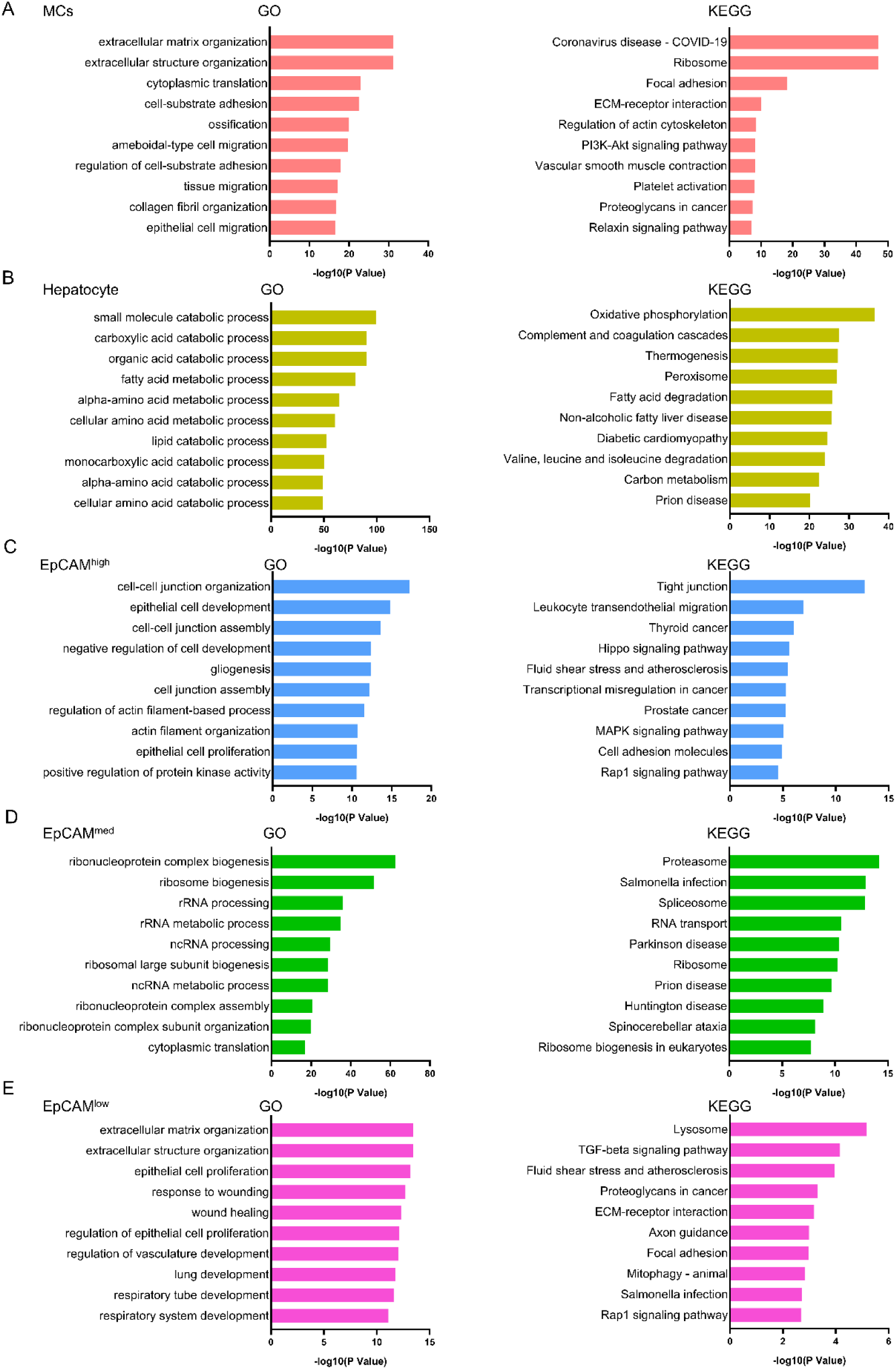
KEGG and GO analyses of liver Tomato^+^ cells of P1 or P14 mice. **(A-E)**. GO and KEGG analyses of various subpopulation using scRNA-seq data.

**Fig. S4.**
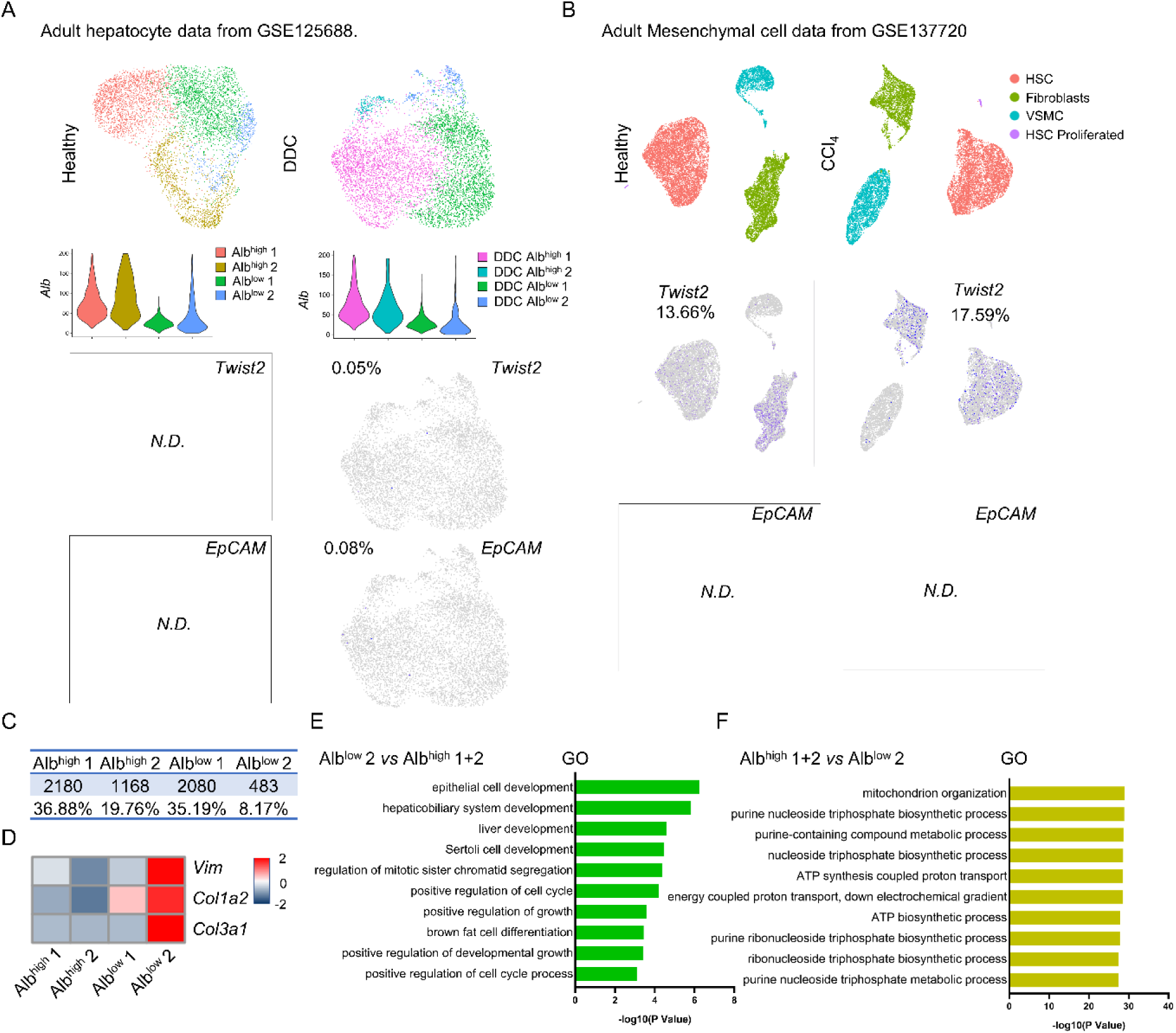
Analysis of the public domain scRNA-seq datasets of hepatocytes and MCs. **(A).** Analyses of hepatocytes from normal mice (based on Alb expression levels) and their expression of *Twist2* and *EpCAM* under normal or injury conditions (GSE125688). DDC (1 week), 3,5-diethoxycarbonyl-1,4-dihydrocollidine. **(B).** t-SNE analysis of liver mesenchymal populations from normal mice and their expression of *Twist2* and *EpCAM* under normal or injury conditions (CCl_4_, 6 weeks) (GSE137720). **(C).** The proportions of various subgroups in the adult mouse liver based on *Alb* expression (GSE125688) (derived from Fig. S4A). **(D).** Heatmap showing the expression of MC markers in hepatocyte subpopulations from adult normal mice (derived from Fig. S4A). **(E).** GO analyses of Alb^low^ 2 hepatocytes compared to Alb^high^ (1+2) hepatocytes. **(F).** GO analyses of Alb^high^ (1+2) compared to Alb^low^ 2 hepatocytes.

**Fig. S5.**
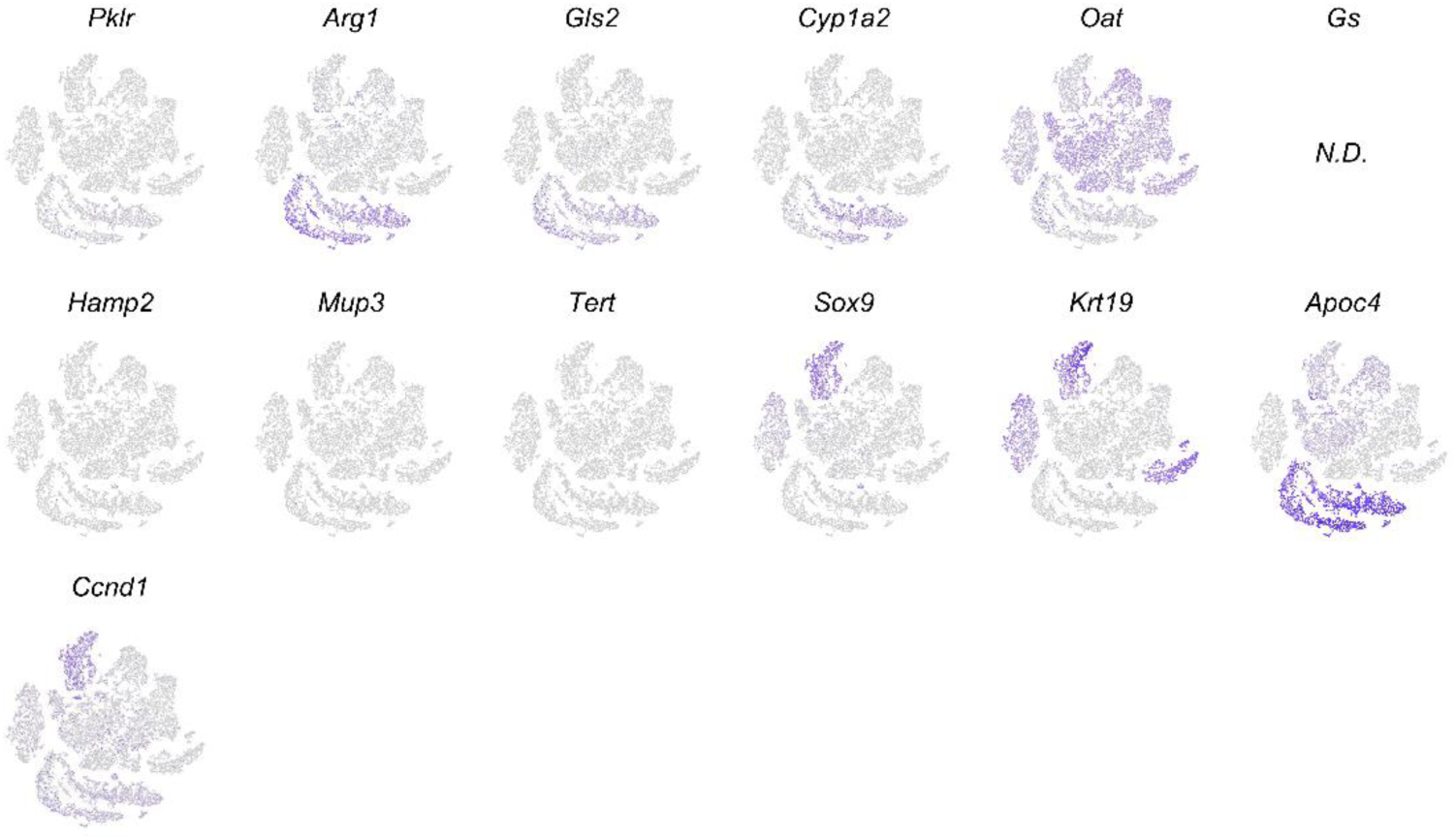
Expression of hepatocyte marker genes in Twist2-lineage liver cells. Analysis of hepatocyte marker gene expression in Twist2-lineage liver cells. These markers were studied to trace hepatocytes in reference 25.

**Fig. S6.**
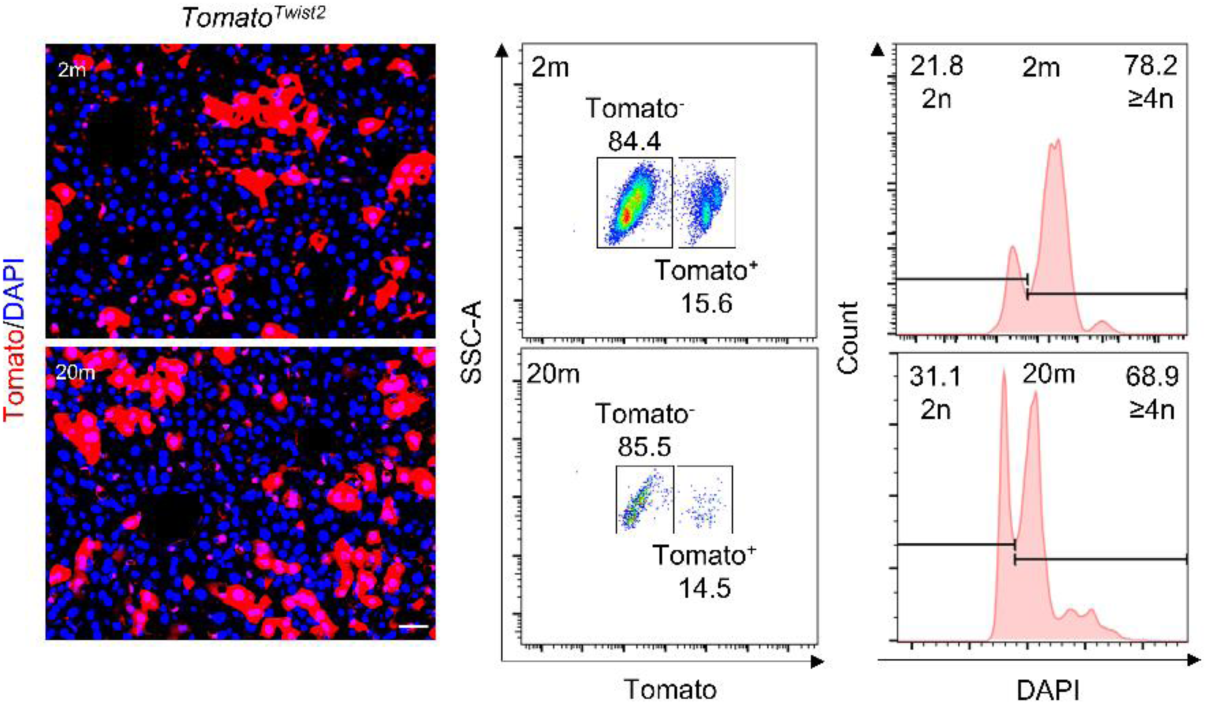
Analysis of Twist2-lineage mesenchymal-hepatocytes in young and old mice. Comparison of the percentage of Tomato^+^ hepatocytes and their polyploid status in 2- and 20-month-old *Tomato^Twist2^* mice by immunostaining and flow cytometry, respectively.

**Fig. S7.**
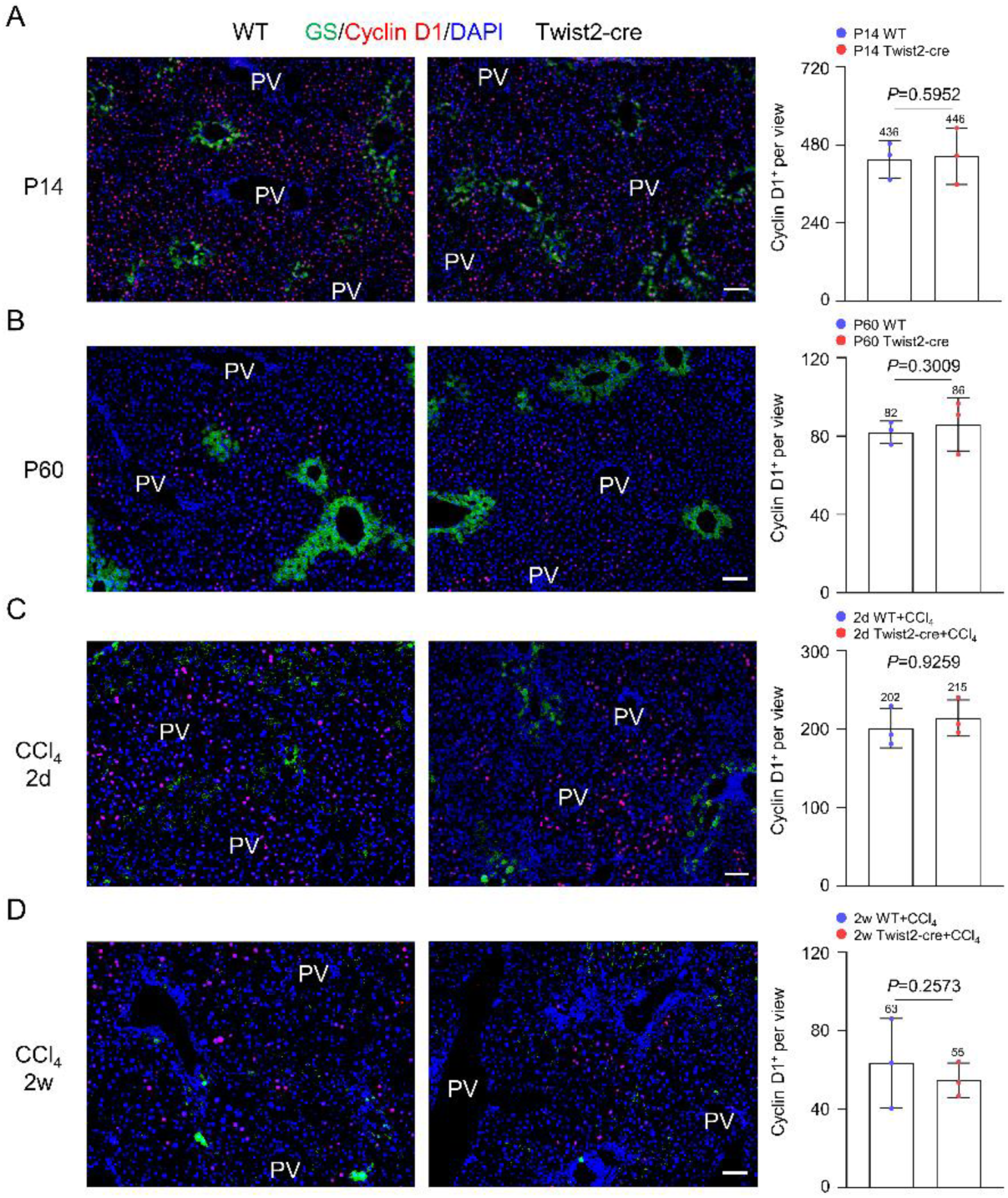
Deletion of one *Twist2* allele did not affect midzone hepatocytes during homeostasis or regeneration. **(A).** Immunostaining results for Cyclin D1 in P14 *Twist2-Cre* and control mice. Right panel: Quantitation data: the numbers of Cyclin D1^+^ cells per view. n=3. Scale bar, 100 µm. Data were shown as mean ±SD. **(B).** Immunostaining results for Cyclin D1 in P60 *Twist2-Cre* and control mice. Right panel: Quantitation data: the numbers of Cyclin D1^+^ cells per view. n=3. Scale bar, 100 µm. Data were shown as mean ±SD. **(C).** Immunostaining results for Cyclin D1 at day 2 post CCl_4_ in adult *Twist2-Cre* and control mice. Right panel: Quantitation data: the numbers of Cyclin D1^+^ cells per view. n=3. Scale bar, 100 µm. Data were shown as mean ±SD. **(D).** Immunostaining results for Cyclin D1 at day 14 post CCl_4_ in adult *Twist2-Cre* and control mice. Right panel: Quantitation data: the numbers of Cyclin D1^+^ cells per view. n=3. Scale bar, 100 µm. Data were shown as mean ±SD.

**Fig. S8.**
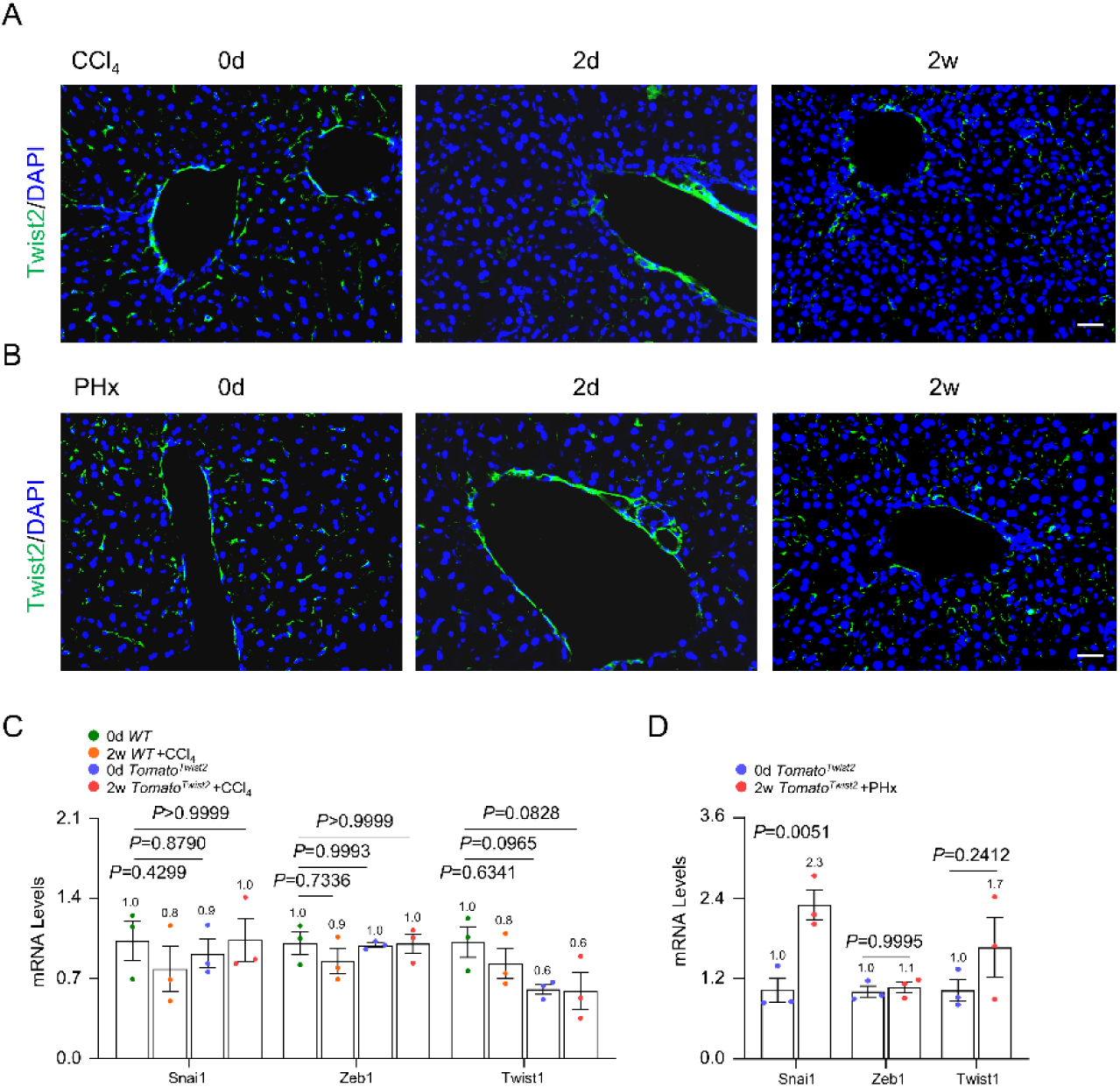
Analysis of expression of Twist2 and MET genes during liver repair. **(A).** Immunostaining results for Twist2 on liver sections of normal adult mice 0 day, 2 days, or 2 weeks after CCl_4_. Scale bar, 50 µm. **(B).** Immunostaining results for Twist2 on liver sections of normal adult mice 0 day, 2 days, or 2 weeks after PHx. Scale bar, 50 µm. **(C).** qPCR results showed that *Snai1*, *Zeb1,* and *Twist1* mRNA levels were not significantly altered in regenerating liver of *Twist2-Cre* mice 2 weeks after CCl_4_ compared to normal mice. n=3. Data were presented as mean ±SD. Two-way ANOVA was applied. *P*-value <0.05 was considered as statistically significant. **(D).** qPCR results showed that *Snai1 mRNA level was increased while Zeb1* and *Twist1* mRNA levels were not significantly altered in regenerating livers of mice 2 weeks after PHx in *Twist2-Cre* mice. n=3. Data were presented as mean ±SD. Two-way ANOVA was applied. *P*-value <0.05 was considered as statistically significant.

**Fig. S9.**
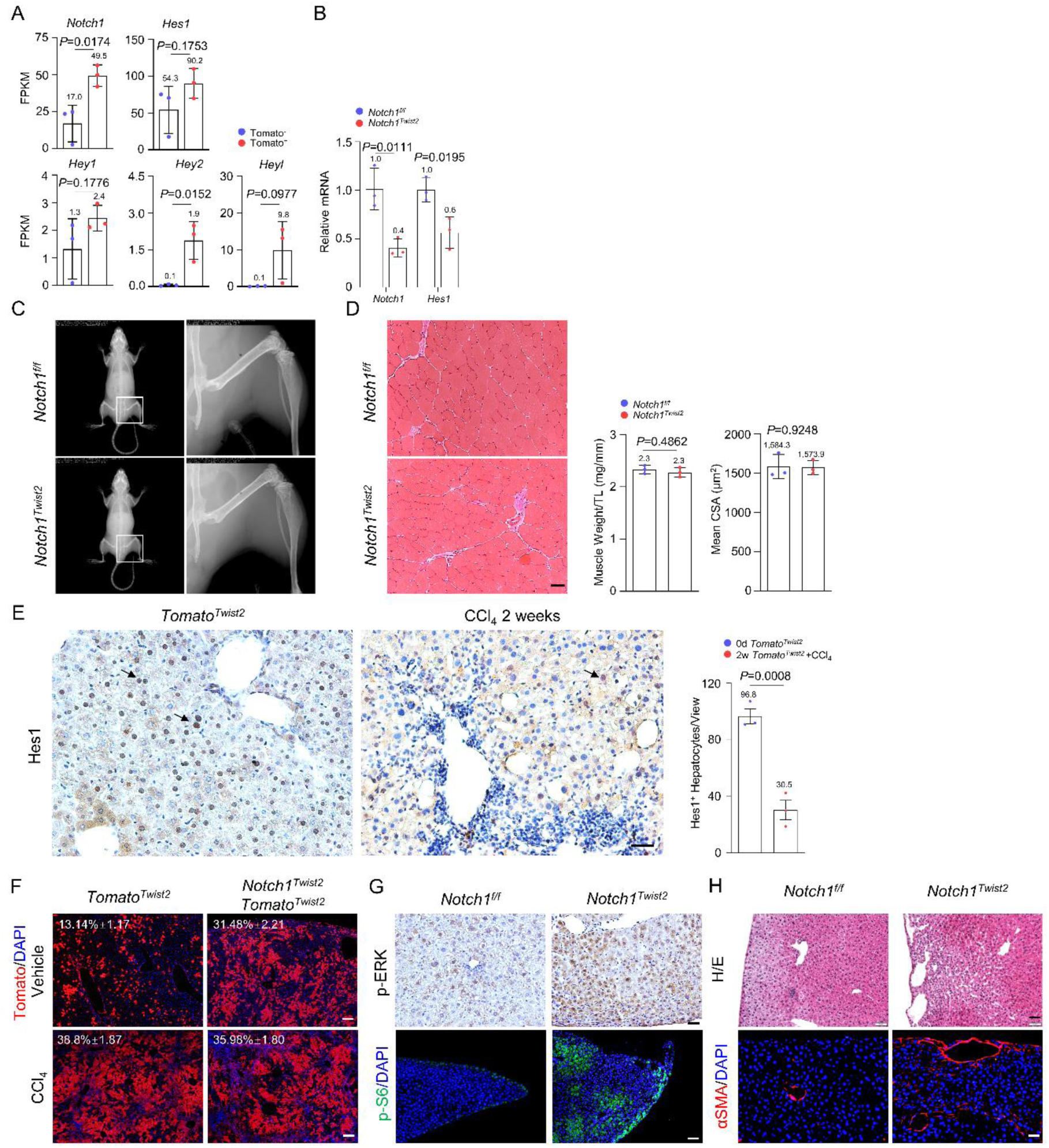
Notch1 limits Twist2-lineage hepatocyte expansion via VEGFR-mTOR signaling. **(A).** FPKM values for major Notch1 pathway genes in Tomato^+^ and Tomato^-^ hepatocytes. n=3, Data were presented as mean ±SD. Unpaired two-tailed Student’s *t*-test was applied. *P*-value <0.05 was considered as statistically significant. **(B).** qPCR results showed a reduction in *Notch1* and *Hes1* in hepatocytes from *Nocth1^Tiwst2^* mice compared to control mice. n=3. Data were presented as mean ±SD. Two-way ANOVA was applied. *P*-value <0.05 was considered as statistically significant. The skeletal structure of adult male *Nocth1^Twsit2^* and control mice. **(C).** The muscle phenotypes of *Nocth1^Twsit2^* and control mice. Right panels: weight to body weight ratio and cross section areas of myofibers. n=3. Data were presented as mean ±SD. Unpaired two-tailed Student’s *t*-test was applied. *P*-value <0.05 was considered as statistically significant. **(D).** Immunostaining results showed reduced expression of Hes1 in regenerating liver samples of normal mice. Scale bar, 50 µm. Right panel: quantitation data. n=3. Data were presented as mean ±SD. Unpaired two-tailed Student’s *t*-test was applied. *P*-value <0.05 was considered as statistically significant. **(E).** Illustrative staining results showing the numbers of Tomato^+^ hepatocytes in *Nocth1^Twist2^*mice and control mice after CCl_4_-induced injury. n=3. Scale bar, 200 µm. Data were shown as mean ±SD. * *P*-value <0.05 between *Nocth1^Twist2^* mice and control mice after CCl_4_-induced injury. **(F).** Analysis of p-Erk and p-S6 on liver sections from adult male *Nocth1^Twist2^* and control mice. Scale bars, 50 µm for p-ERK and 100 µm for p-S6. **(G).** The *Nocth1^Twist2^* mice showed an increase in blood vessels in the liver, which were stained positive for αSMA. Scale bar, 50 µm.

**Fig. S10.**
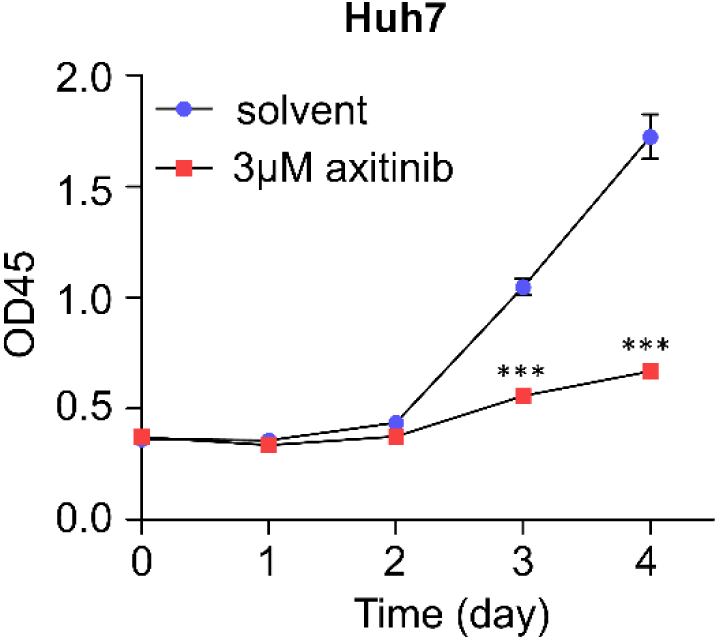
Axitinib inhibits proliferation of Huh7 cells. CCK-8 assays were performed to evaluate the effect of Axitinib on the proliferation of hepatocellular carcinoma cell line Huh7. n=5. Data were presented as mean ±SD. Two-way ANOVA was applied. Statistical significance is displayed as *P*<0.001 (***).

